# Olfactory cortical outputs recruit and shape distinct brain-wide spatiotemporal networks

**DOI:** 10.1101/2024.07.19.604242

**Authors:** Teng Ma, Xunda Wang, Xuehong Lin, Junjian Wen, Linshan Xie, Pek-Lan Khong, Peng Cao, Ed X. Wu, Alex T. L. Leong

**Author notes:** Correspondence should be addressed to Alex T. L. Leong, Ph.D.: Laboratory of Biomedical Imaging and Signal Processing, Department of Electrical and Electronic Engineering, The University of Hong Kong, Pokfulam, Hong Kong, Hong Kong SAR, China.

## Abstract

Odor information is transmitted from the olfactory bulb to several primary olfactory cortical regions in parallel, including the anterior olfactory nucleus (AON) and piriform cortex (Pir). However, the specific roles of the olfactory bulb and cortical outputs in wider interactions with other interconnected regions throughout the brain remain unclear due to the lack of suitable in vivo techniques. Furthermore, emerging associations between olfactory-related dysfunctions and neurological disorders underscore the need for examining olfactory networks at the systems level. Using optogenetics, fMRI, and computational modeling, we interrogated the spatiotemporal properties of brain-wide neural interactions in olfactory networks. We observed distinct downstream recruitment patterns. Specifically, stimulation of excitatory projection neurons in OB predominantly activates primary olfactory network regions, while stimulation of OB afferents in AON and Pir primarily orthodromically activates hippocampal/striatal and limbic networks, respectively. Temporally, repeated OB or AON stimulation diminishes neural activity propagation brain-wide in contrast to Pir stimulation. Dynamic causal modeling analysis reveals a robust inhibitory effect of AON outputs on striatal and limbic network regions. In addition, experiments in aged rat models show decreased brain-wide activation following OB stimulation, particularly in the primary olfactory and limbic networks. Modeling analysis identifies a dysfunctional AON to Pir connection, indicating the impairment of this primary olfactory cortical circuit that disrupts the downstream long-range propagation. Our study delineates the spatiotemporal properties of olfactory neural activity propagation in brain-wide networks for the first time and distinguishes the roles of primary olfactory cortical, AON and Pir, outputs in shaping neural interactions at the systems level.

## Introduction

The olfactory system plays a crucial role in driving behavioral responses that are essential for daily functions. This system facilitates social behaviors by integrating olfactory stimuli with other sensory stimuli and prior experiences^1,2^. Olfactory systems can discriminate among hundreds to thousands of different odorants given the diversity of odorant receptors that specifically bind to the corresponding odorant molecules^3,4^. While olfaction may be more critical for lower-level mammalian species, it remains important for higher primates, as olfactory stimuli can influence emotions, moods, and behaviors^5^. One unique feature of the anatomical organization of the olfactory system is the absence of thalamic processing, indicating that the brain networks pivotal for olfactory processing are distinctly different from other sensory systems. Odor information detected in the olfactory epithelium is first processed at the olfactory bulb (OB), before being routed in parallel to several primary olfactory cortical regions, including the anterior olfactory nucleus (AON) and piriform cortex (Pir). Present studies have primarily examined anatomical projections and targets of local olfactory microcircuits, such as the OB, AON and Pir, including the synaptic interactions between these three connected regions^6–8^. However, the neural interplay between small numbers of anatomically connected regions (i.e., local micro-circuits) is likely insufficient to describe the dynamic properties of information processing that underlie specific brain functions^9–11^. Recent views have advocated for examining and/or modeling polysynaptic interactions across complex networks spanning multiple circuits distributed throughout the brain^11,12^. Moreover, the associations between olfactory-related dysfunctions and neurological disorders such as aging^13^, neurodegenerative diseases^14^, and recently COVID-19^15–17^, highlight the need for examining the olfactory network at the systems level.

Interrogating brain-wide, long-range olfactory networks and their properties require a large-view brain-wide imaging technique that is sensitive to neural activity. However, mapping the brain-wide extent of olfactory regions via traditional functional MRI (fMRI) mapping approaches is technically challenging due to prominent olfactory response habituation^18,19^. Further, the need to present a multitude of odor combinations also makes it challenging to efficiently and reliably interrogate long-range olfactory networks and their properties. Previous attempts to study brain-wide regions involved in olfactory processing in rodents^20,21^ and humans^22,23^ using task-based fMRI have only identified some known primary olfactory regions, such as the OB, AON, Pir, olfactory tubercle (Tu), and amygdala (Amg). There is a significant gap in the visualization of long-range olfactory networks and the characterization of their functional properties. Notably, the specific roles of the bulbar and AON or Pir cortical outputs in wider interactions with other interconnected regions throughout the brain in olfactory processing remain unclear. In addition, several higher-order regions suspected to mediate more complex olfactory processing remain unidentified and/or to be characterized^5,24,25^. Thus, we designed alternative neuromodulation strategies that can stimulate olfactory-specific neurons in the OB, AON, and Pir with fMRI to characterize brain-wide downstream activation targets and the dynamic properties of long-range olfactory networks. We also aim to employ dynamic causal modeling (DCM)^26^, a computational framework that can be used to infer causal interactions among different regions in the olfactory network under different stimulation conditions.

In this study, we deployed optogenetic fMRI to achieve reversible, millisecond precision, and cell-type specific stimulation^12,27–31^ of excitatory projection neurons in OB (i.e., mitral and tufted cells), and the olfactory-specific neurons of AON and Pir with computational modeling to investigate the dynamics of large-scale olfactory spatiotemporal networks. Our findings revealed distinct recruitment of downstream targets by olfactory-specific neural populations in AON and Pir. AON-driven neural activities strongly activated hippocampal and striatal networks, while Pir-driven activities preferentially recruited the limbic network. Repeated excitations of AON or OB across multiple fMRI sessions decreased brain-wide activations, while Pir excitations did not alter orthodromic neural activity propagation in long-range olfactory networks. DCM of optogenetic fMRI data showed consistent negative effective connectivity from AON to various downstream targets in the striatal and limbic networks, indicating a robust inhibitory effect of AON outputs on neural activity propagation in long-range olfactory networks. Further, our systematic examination of long-range olfactory networks in an aged rat model demonstrated an overall decrease in brain-wide activations upon OB excitations, particularly in the primary olfactory and limbic networks. DCM showed that the effective connectivity from AON to Pir switched from positive (healthy) to negative (aged), indicating impairment of this cortical circuit that disrupted neural activity propagation to primary olfactory and limbic networks. This study provides the initial delineation of the spatiotemporal properties of olfactory neural activity propagation in brain-wide networks. We show that primary olfactory cortices (AON and Pir) recruit distinct higher-order brain networks. We also uncover the role of the inhibitory effect of AON and the excitatory effect of Pir outputs in shaping the dynamic properties of neural activity propagation in long-range olfactory networks.

## Results

### Olfactory bulb and primary olfactory cortices recruit distinct brain-wide long-range networks

To identify activated neural networks of the olfactory system, we fused Ca^2+^/calmodulin-dependent protein kinase II alpha (CaMKIIα)-dependent ChR2(H134R) with mCherry to transfect excitatory projection neurons in OB of healthy adult Sprague Dawley rats. Colocalization of mCherry and CaMKIIα confirmed the specific expression of CaMKIIα in mitral and tufted cells within the mitral cell layer and the external plexiform layer of OB (**Fig. 1A**). Optogenetic stimulation was delivered at 1 Hz (10% duty cycle, 40 mW/mm^2^), 5, 10, 20 and 40 Hz (30% duty cycle, 40 mW/mm^2^) in a block-design paradigm covering the frequency range of delta (1 Hz, low frequency) to gamma (40 Hz, high frequency) oscillations. Different frequencies were presented in a randomized manner within each fMRI session (**Fig. 1B**). In particular, 1 Hz is closest to the most biologically relevant frequency during olfaction due to the intrinsic respiration-related oscillatory properties (i.e., 1.5 Hz) within the olfactory system^32^.

**Figure 1.**
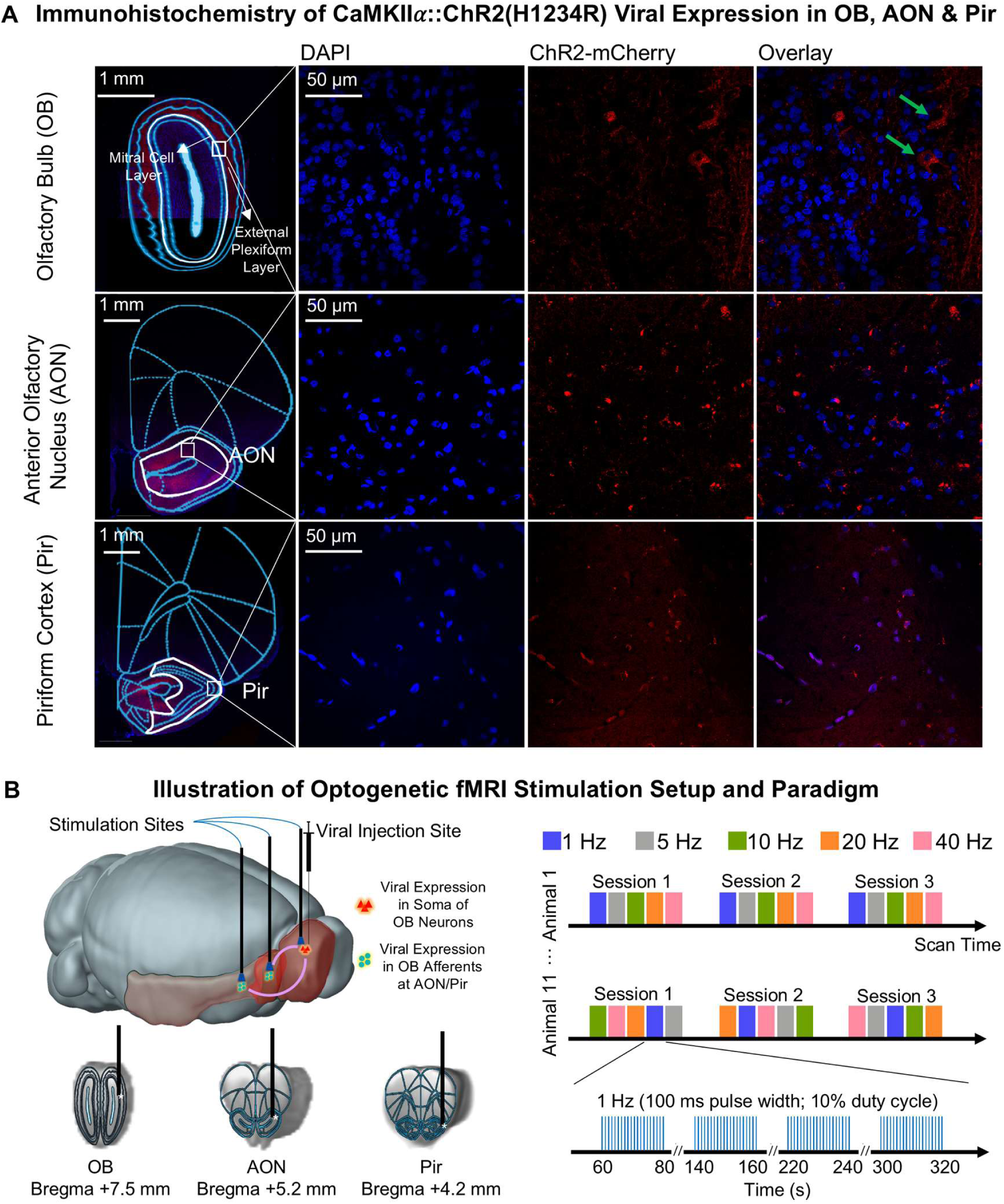
Histological characterization of ChR2::CaMKIIα viral expression in the olfactory bulb (OB) excitatory neurons and OB afferents at the anterior olfactory nucleus (AON) and the piriform cortex (Pir), and optogenetic fMRI stimulation setup. **(A)** Confocal images, 10x magnification (Left) and 40x magnification (Right), of ChR2-mCherry expression in histological slices covering OB, AON and Pir. Overlay of images co-stained for the nuclear marker DAPI and mCherry in OB (Top) revealed ChR2 expression in the soma of the mitral cell layer & external plexiform layer in OB (indicated by green arrows). Meanwhile, ChR2 expressions were found in OB afferents at AON and Pir (Middle & Bottom), indicated by minimal colocalization between mCherry and DAPI. **(B)** Illustration of three optogenetic stimulation targets, namely the OB excitatory neurons, and OB afferents in AON and Pir (Left), which were conducted in separate animal groups. T2-weighted anatomical MRI image showing the location of the implanted optical fiber (asterisk: stimulation target). Optogenetic fMRI block-designed stimulation paradigm (Right). Five different frequencies (1, 5, 10, 20, and 40 Hz) were presented in a pseudorandomized order within each session. We repeated the stimulations for a total of three sessions in each animal fMRI experiment. All frequencies were presented at a 30% duty cycle except for 1 Hz, which was presented at a 10% duty cycle.

First, we stimulated the OB, the first stage of olfactory information processing in the central nervous system. We detected robust BOLD fMRI activation bilaterally within and beyond primary olfactory networks during 1 Hz stimulation in ipsilateral OB excitatory (**Fig. 2A**). We identified robust and widespread BOLD responses across both hemispheres in known primary olfactory (AON, Pir, tenia tecta, TT, entorhinal cortex, Ent and olfactory tubercle, Tu), limbic (cingulate, Cg, orbitofrontal, OFC and insular, Ins, cortices, and amygdala, Amg), hippocampal (ventral hippocampus, vHP), striatal (nucleus accumbens, NAc and ventral caudate putamen, vCPu), and sensorimotor (motor, MC, somatosensory, S1, auditory, AC and visual, V1, cortices) regions (**Fig. 2A**). These activations indicate the presence of interactions between the primary olfactory network and other networks associated with high-order functions beyond olfactory processing, such as cognition, reward, and cross-modal sensory processing.

**Figure 2.**
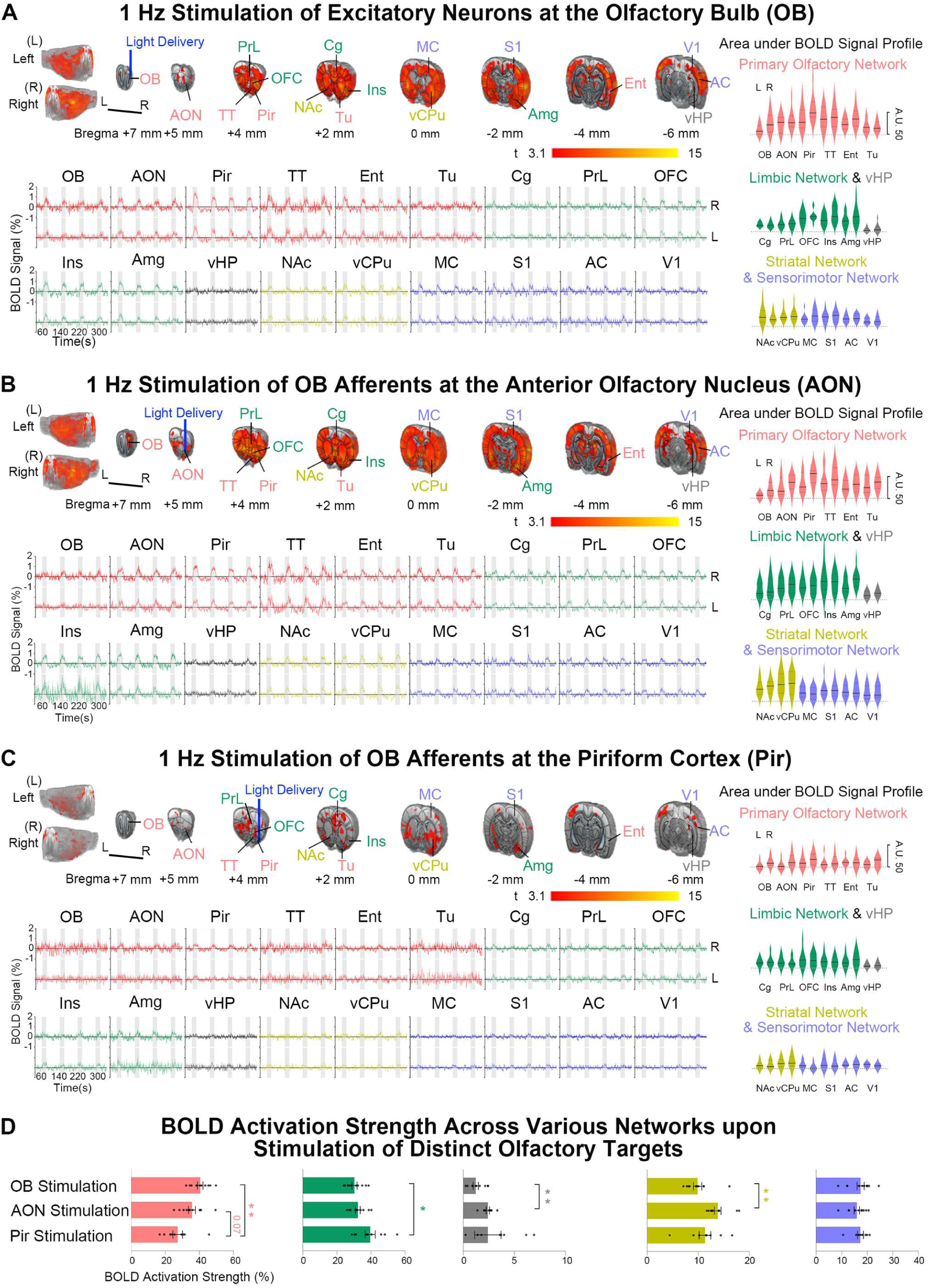
Optogenetic excitation of OB excitatory neurons and OB afferents at AON and Pir recruit distinct brain-wide long-range olfactory networks. Averaged BOLD activation maps and signal profiles (Left) upon 1 Hz stimulation of (**A**) OB excitatory neurons, (**B**) OB afferents in AON, and (**C**) OB afferents in Pir during the first fMRI session (n = 11 for fMRI experiments in A & B; n = 9 for C; error bars indicate ± SEM), and the corresponding area under the BOLD signal profiles (AUC) for each atlas-defined region-of-interests (ROIs; Right). Note that the BOLD activation maps displayed in A-C were generated as follows. First, uncorrected one-sample t-tests were conducted with a threshold of P < 0.001, corresponding to t > 3.1. 2). These thresholded activation maps were further corrected using nonparametric inference with threshold-free cluster enhancement multiple comparison correction of family-wise error rate (TFCE-FWE, P < 0.05). (**D**) Summary of BOLD activation strength to quantify the extent of various brain networks engaged upon respective optogenetic stimulation of the three different primary olfactory stimulation targets. BOLD activation strength was computed as the ratio of AUC for each ROI over AUC of all ROIs (individual animal data points are shown; error bars indicate ± SEM; Tukey’s multiple comparisons test; \*\**P* < 0.01, and \**P* < 0.05). OB-driven neural activities mainly activated the primary olfactory network (AON, Pir, TT, Ent and Tu). AON-driven neural activities were found to preferentially recruit the striatal (NAc and vCPu) and hippocampal networks (vHP), while Pir-driven activities strongly targeted the limbic network (Cg, PrL, OFC, Ins and Amg). *Abbreviations. Regions in primary olfactory network: olfactory bulb (OB), anterior olfactory nucleus (AON), piriform cortex (Pir), tenia tecta (TT), entorhinal cortex (Ent), olfactory tubercle (Tu); limbic network: cingulate (Cg), prelimbic cortex (PrL), orbitofrontal cortex (OFC), insular (Ins), amygdala (Amg); hippocampal network: ventral hippocampus (vHP); striatal network: nucleus accumbens (NAc), ventral caudate putamen (vCPu); sensorimotor network: motor (MC), somatosensory (S1), auditory (AC), visual (V1) cortex.*

Although the first stage of olfactory processing occurs at the OB, primary olfactory cortices (e.g., AON and Pir), which receive parallel direct input from the OB, are also critical for olfaction, as they participate in olfactory memory functions and learning in odor representations within local micro-circuits^33,34^ and interactions with the hippocampus^33,35^. As such, we sought to directly examine AON/Pir in our olfactory brain-wide neural networks. We note that direct optogenetic targeting of AON/Pir excitatory neurons may target additional neuronal populations, as not all excitatory neuronal populations in AON and Pir receive olfactory inputs from OB^36,37^. Here, we designed an optogenetic stimulation protocol based on projection targeting (**Supplementary Fig. 1**), whereby the OB afferent terminals were stimulated at AON/Pir, thereby enabling selective stimulation of neurons in AON/Pir that exclusively receive olfactory inputs. Upon 1 Hz stimulation of OB afferents at AON, we observed robust widespread BOLD activations in bilateral primary olfactory, limbic, striatal, sensorimotor cortical and hippocampal regions (**Fig. 2B**), similar to the spatial patterns in our findings for OB stimulation. Meanwhile, 1 Hz optogenetic stimulation of OB afferents at Pir also evoked bilateral BOLD responses in primary olfactory, limbic, and striatal regions albeit with significantly weaker and sparser BOLD activations in bilateral limbic, striatal, and sensorimotor cortical regions (**Fig. 2C**). Note that the EPI images from a representative animal that underwent an optogenetic fMRI experiment showed no appreciable signal dropout in olfactory regions near air-tissue interfaces such as OB, AON, Pir and Amg (**Supplementary Fig. 2**). Hence, subsequent quantitative analyses involving these regions were unlikely to be affected.

In addition to the qualitative description of BOLD activation distributions, we further sought to quantify the strength of neural activity at each downstream activation target upon stimulation of the three targets (i.e., OB, AON and Pir). We first calculated the area under the BOLD signal profile (AUC) during the four 20-s 1 Hz optogenetic stimulation blocks for each downstream activation target (**Fig. 2A-C**). However, direct comparisons between AUCs of the three separate optogenetic fMRI experiments are erroneous given the differences in absolute BOLD signal amplitudes, which are likely caused by differences in ChR2 transfection volumes in OB excitatory neurons and OB afferents at AON/Pir, and subsequently the population size of neurons/axonal terminals stimulated. The vast majority of OB neurons are interneurons and not projection neurons that terminate in AON/Pir^38^. Hence, we normalized the AUCs to the sum of BOLD activations of their respective optogenetic fMRI experiments (i.e., defined as BOLD activation strength here; **Fig. 2D**). To distinguish the brain networks that are predominantly recruited by AON- or Pir-driven neural activities compared to OB stimulation, we found that the bulk of OB-driven neural activities were concentrated within the primary olfactory network (i.e., OB, AON, Pir, TT, Ent and Tu; 40.5 ± 1.6 % vs. 27.2 ± 2.7 %, *P* < 0.01 for OB vs. Pir; and 40.5 ± 1.6 % vs. 35.6 ± 2.0 %, *P* < 0.1 for OB vs. AON). The strongest BOLD activations within the primary olfactory network (i.e., AON, Pir, TT, Ent, Tu) for OB stimulation experiments aligned with the direct anatomical projections from OB to these targets^38^, indicating the successful excitation of OB projection neurons. Interestingly, we discovered that AON-driven activities preferentially recruited the striatal (i.e., NAc and vCPu; 13.8 ± 0.6 % vs. 9.8 ± 0.8 %, *P* < 0.01 for AON vs. OB) and hippocampal networks (2.4 ± 0.1 % vs. 1.2 ± 0.2 %, *P* < 0.01 for AON vs. OB), while Pir-driven activities strongly targeted the limbic network (i.e., Cg, PrL, OFC, Ins and Amg; 39.5 ± 2.8 % vs. 30.3 ± 1.4 %, *P* < 0.05 for Pir vs. OB). Meanwhile, all three stimulated regions showed relatively equal recruitment of sensorimotor networks. The specificity afforded by perturbing the OB afferents at AON and Pir indicated that these olfactory inputs recruit distinct downstream targets in primary olfactory, limbic, hippocampal and striatal networks, despite sharing extensive overlaps in anatomical projections^25,38–40^.

We also examined BOLD activations upon optogenetic stimulation of the three primary olfactory regions (i.e., OB, AON and Pir) at various frequencies. We observed robust bilateral activations during 5 Hz stimulation of OB excitatory neurons (**Supplementary Fig. 3A**) and OB afferents in AON (**Supplementary Fig. 4A**), although the activations at the contralateral hemisphere were relatively weaker. The subsequent increase in stimulation frequencies to 10 Hz, 20 Hz and 40 Hz resulted in robust activations that were predominantly localized in the ipsilateral primary olfactory and limbic networks (**Supplementary Figs. 3B-D & 4B-D**). Meanwhile, BOLD activations were only detected in ipsilateral primary olfactory, limbic, and striatal networks when higher frequency optogenetic stimulation was presented to OB afferents at Pir (**Supplementary Fig. 5**). To examine long-range olfactory networks, we focused on the 1 Hz optogenetic stimulation experiments.

Altogether, our approach to stimulating the three primary olfactory regions (i.e., OB, AON and Pir) provides valuable insights into the networks that are pivotal for olfactory processing. Notably, the distinction in brain networks that are recruited by olfactory-specific neural populations in AON vs. Pir such as hippocampal and striatal vs. limbic reveal functional insight into the AON and Pir in olfactory processing.

### AON and Pir circuit outputs shape the dynamic properties of long-range olfactory networks

The distinct spatiotemporal propagation properties and the recruitment of different long-range olfactory-related networks suggest varied modulatory roles for OB, AON and Pir when interacting with their downstream targets. Here, we first examined the dynamics of BOLD activations across multiple 1 Hz stimulation sessions. We observed distinct changes to the spatiotemporal properties of BOLD fMRI activations across the three optogenetic fMRI sessions. Overall, the brain-wide activations at primary olfactory, limbic, striatal, and sensorimotor cortical networks in our first fMRI session showed an appreciable decreasing trend in subsequent stimulation sessions. For OB 1 Hz stimulation in the 2^nd^ fMRI session, there was a decrease in the BOLD activations at most of the downstream targets across multiple networks. These include the ipsilateral OB (i.e., the stimulation site), contralateral TT, and bilateral AON, Pir, Ent and Tu of the primary olfactory network; bilateral OFC, Ins and Amg of the limbic network; bilateral NAc and ipsilateral vCPu of the striatal network; and bilateral MC and ipsilateral AC of the sensorimotor network (**Fig. 3A** and **Supplementary Fig. 6A**). BOLD activations during the 3^rd^ fMRI session also showed decreased activations (i.e., 3^rd^ vs 1^st^ fMRI session; **Fig. 3A** and **Supplementary Fig. 6A**). Comparatively, we observed a dramatic decrease in activations upon 1 Hz stimulation of OB afferents at AON during the later fMRI sessions. We found diminished or even absent bilateral activations in downstream targets. Specifically, activations evoked by AON stimulation showed a significant decrease in the 2^nd^ and 3^rd^ sessions at all regions except contralateral OB (**Fig. 3B** and **Supplementary Fig. 6B**). Notably, activations at bilateral vHP also significantly decreased despite having the weakest activation among downstream targets during the 1^st^ fMRI session. However, the activations upon 1 Hz stimulation of OB afferent inputs at Pir remained relatively stable across fMRI sessions in the primary olfactory, limbic, striatal and sensory cortical networks except for a slight decrease in Amg (**Fig. 3C** and **Supplementary Fig. 6C**).

**Figure 3.**
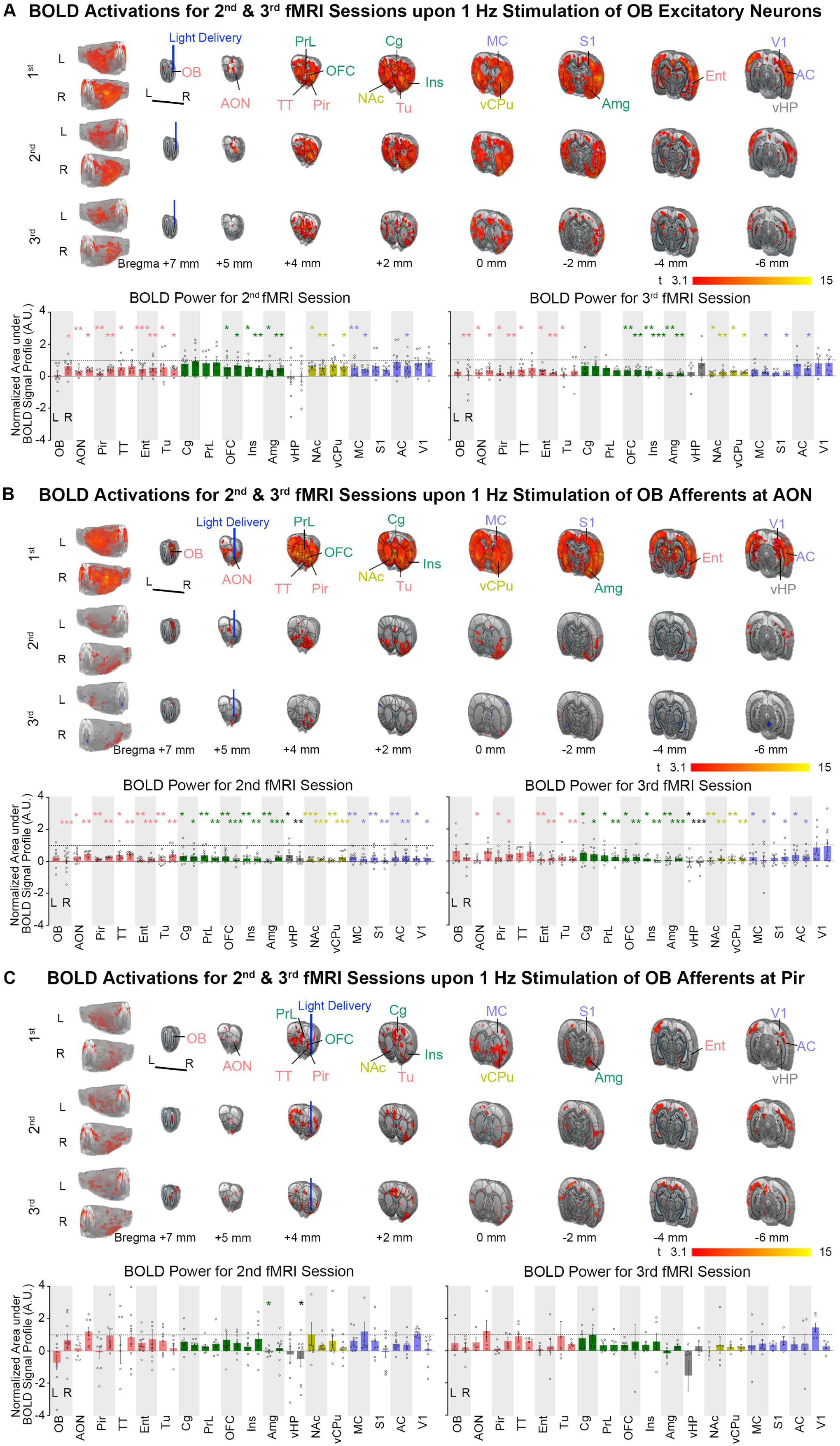
Repetitive optogenetic excitation of OB excitatory neurons and OB afferents at AON and Pir reveal varied neural activity adaptation properties in olfactory networks. Averaged BOLD activation maps (Top) upon 1 Hz stimulation of (**A**) OB excitatory neurons, (**B**) OB afferents in AON, and (**C**) OB afferents in Pir during the first, second (n = 11 for fMRI experiments in A & B; n = 9 for C; t > 3.1 corresponding to *P* < 0.001) and third (n = 8 for fMRI experiments in A; n = 9 for B; n = 6 for C; t > 3.1 corresponding to *P* < 0.001) fMRI sessions, respectively. We found a dramatic decrease or even absent bilateral activations in long-range olfactory networks upon stimulation of OB excitatory neurons or OB afferents at AON, but not Pir, indicating strong neural activity adaptation mediated by AON-driven neural activity to downstream targets. Note that similar to Fig. 2, the BOLD activation maps displayed in A-C were further corrected for multiple comparisons with threshold-free cluster enhancement with family-wise error rate (TFCE-FWE) at *P* < 0.05, except for that the maps during the third session in B and C were generated directly after one-sample t-test. The extent of neural activity adaptation in brain-wide olfactory networks was quantified with the true BOLD power (AUC of the BOLD signal profiles) for each atlas-defined ROI (Bottom; individual animal data points are shown; error bars indicate ± SEM; Dunnett’s multiple comparisons test of BOLD power during 2^nd^ vs. 1^st^ fMRI session, 3^rd^ vs. 1^st^ session; \*\*\**P* < 0.001; \*\**P* < 0.01; \**P* < 0.05). Note that the AUC for each ROI was normalized to their respective counterpart in the first fMRI session for display.

We then sought to identify the driver of these distinct influences of OB, AON and Pir stimulation on the dynamic response properties of long-range olfactory-related networks. Conventional analyses of BOLD fMRI data such as general linear model (GLM) and comparisons of the energy of BOLD signal profiles are insufficient and unable to infer the causal influences of OB-, AON- and Pir-driven neural activity on their downstream targets. Here, we applied DCM^26^ on our optogenetic fMRI data to parametrize the downstream effects on selected targets in the right brain hemisphere (i.e., the same hemispheric side to the optogenetic stimulation site). Specifically, we used DCM to computationally model the effective connectivity strength of selected downstream connections in OB, AON and Pir stimulation data. We defined a six-node brain network and its connections that encompassed regions from primary olfactory (OB, AON, Pir and Ent), striatal (vCPu) and limbic (Amg) networks (**Fig. 4A**). These connections were primarily based on the well-defined anatomical feedforward and feedback projections involving the three major primary olfactory regions (i.e., OB, AON and Pir). For simplicity, we did not model other downstream regions to avoid data overfitting issues^41^. Further, we only modeled the optogenetic fMRI data from the 1^st^ session, because BOLD signal profiles for the 2^nd^ and 3^rd^ fMRI sessions had lower SNR due to decreased and in some cases absent activations in long-range olfactory networks. Overall, we observed that the connectivities were primarily positive (i.e., excitatory effects) between the nodes in the primary olfactory network (i.e., OB, AON and Pir; **Fig. 4B**) for all three different stimulations, corroborating documented reciprocal glutamatergic excitatory projections between these three regions^25,39,42,43^. Notably, we observed a significant negative AON and positive Pir connectivity to downstream targets (i.e., Ent, vCPu and Amg; **Fig. 4B**). Quantitative comparisons of the effective connectivity strengths from AON/Pir to Ent, vCPu and Amg confirmed the robust negative AON vs. positive Pir connectivity to downstream targets (*AON-Ent vs. Pir-Ent*: −0.21 ± 0.14 vs. 0.39 ± 0.08 Hz, *P* < 0.05 for OB stimulation and −0.34 ± 0.09 vs. 0.35 ± 0.06 Hz, *P* < 0.01 for AON stimulation; *AON-vCPu vs. Pir-vCPu*: −0.03 ± 0.06 vs. 0.31 ± 0.10 Hz, *P* = 0.053 for OB stimulation, −0.37 ± 0.08 vs. 0.40 ± 0.06 Hz, *P* < 0.01 for AON stimulation and −0.14 ± 0.06 vs. 0.28 ± 0.07 Hz, *P* < 0.05 for Pir stimulation; *AON-Amg vs. Pir-Amg*: −0.10 ± 0.10 vs. 0.26 ± 0.04 Hz, *P* < 0.05 for OB stimulation, −0.25 ± 0.12 vs. 0.33 ± 0.06 Hz, *P* < 0.05 for AON stimulation and −0.16 ± 0.06 vs 0.30 ± 0.06 Hz, *P* < 0.05 for Pir stimulation; **Fig. 4C** and **Supplementary Fig. 7A-C**). These results demonstrate distinct inhibitory effect of AON and excitatory effect of Pir outputs in long-range olfactory networks. We speculate that the significant decrease in BOLD activations during repeated OB or AON stimulations, but not during Pir stimulations, arises from the strong inhibitory effect of AON outputs to various targets beyond the local olfactory microcircuits (i.e., OB, AON and Pir). Taken together, our results indicate that the outputs of primary olfactory cortices, AON and Pir, shape the dynamic properties of neural activity propagation in long-range olfactory networks.

**Figure 4.**
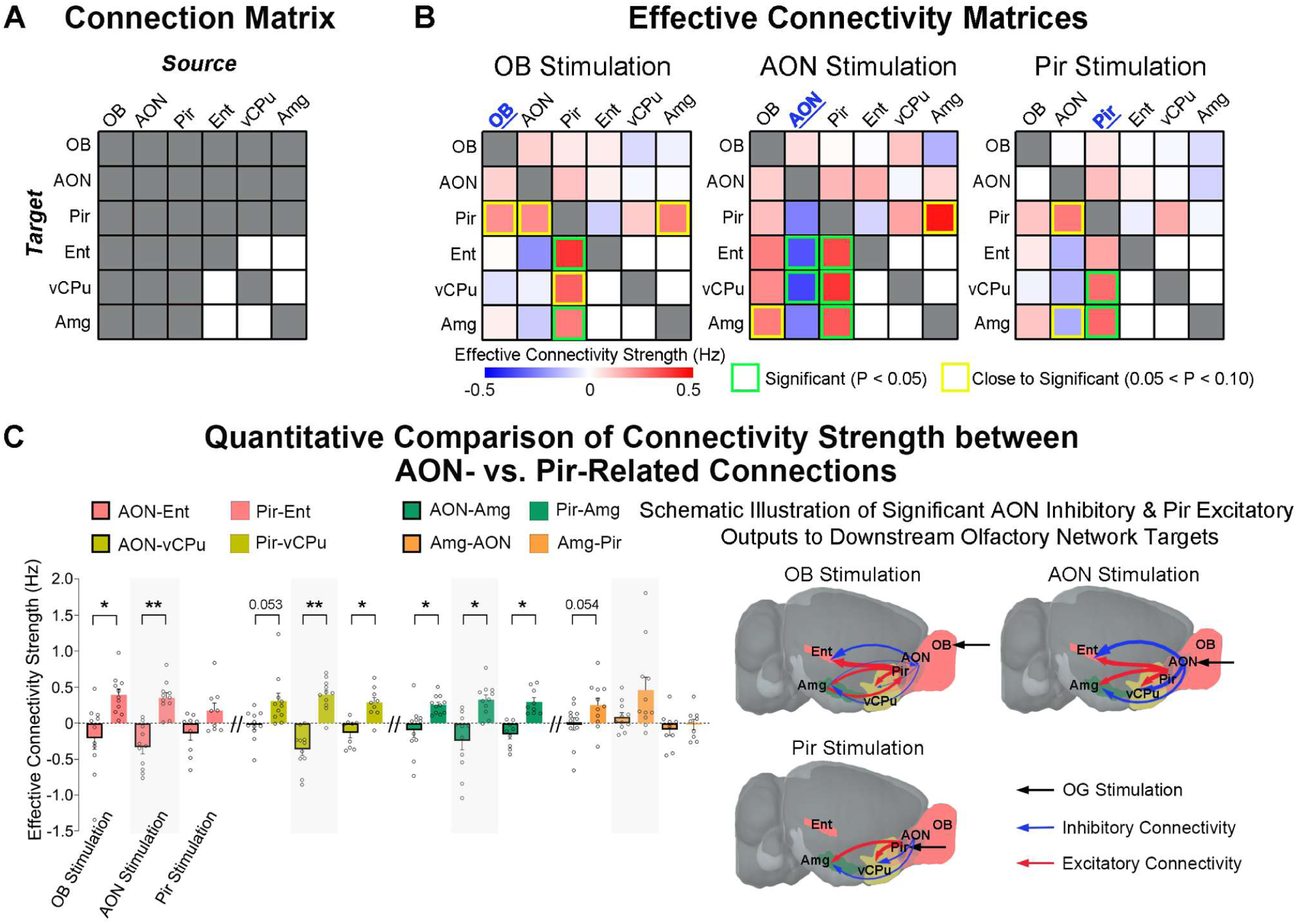
Inhibitory effect of AON outputs and excitatory effect of Pir outputs to downstream targets in long-range olfactory networks. (**A**) Graphical representation of matrix A which defines the a priori anatomical connections of the olfactory network model utilized in dynamic causal modeling (DCM). (**B**) Effective connectivity matrices of the olfactory network model upon stimulation of OB excitatory neurons and their afferents at AON and Pir. (**C**) Quantitative summary of the significant differences in connectivity strength of downstream connections identified from the statistical comparison of matrices in B (Left; n = 11 for OB and AON stimulations; n = 9 for Pir stimulation; individual animal data points are shown; error bars indicate ± SEM; paired t-test with FDR correction; \*\**P* < 0.01, and \**P* < 0.05) and the corresponding graphical representation of inhibitory effect of AON and excitatory effect of Pir outputs to downstream targets (Right). We identified robust AON negative vs. Pir positive connectivity to downstream targets such as Ent, vCPu and Amg, indicating distinct inhibitory vs. excitatory effect of the two primary olfactory cortical outputs that can shape the dynamic properties (i.e., BOLD activations across repeated stimulation sessions) of long-range olfactory networks.

### Repeated AON stimulations diminish orthodromic neural activity propagation

We recognize that our optogenetic stimulation protocols, particularly the stimulations of ChR2-transfected OB afferents at AON and Pir compared to the stimulation of OB excitatory neurons (**Supplementary Fig. 1**), can be sensitive to antidromic-driven neural activities due to activation of the axon terminals instead of the soma^44,45^. To verify the neural activities underlying our fMRI findings and delineate features of neural activity propagation, we conducted local field potential (LFP) recordings at seven regions in the right brain hemisphere. These regions include the ipsilateral OB, AON, Pir, vCPu, V1, Amg, vHP and dHP. The stimulation paradigm (i.e., block-designed and three recording sessions for each animal) matches that from the fMRI experiments. The averaged four 60-s-block LFP traces (**Fig. 5A-C**) and the subsequent power analysis of these LFP traces for the first recording session (**Fig. 6A-C**) largely concurred with the magnitude and temporal profiles of BOLD fMRI activations (**Fig. 2A-C**), with evoked LFPs observed in all recorded regions during the first experimental session. Parallel to our fMRI findings, the strength of neural activities, which were quantified through LFP power analysis (i.e., area under absolute LFP trace normalized to 30-s baseline LFP power), was also decreased during repetitive 1 Hz stimulation of OB or OB afferents at AON but was relatively unchanged under Pir stimulation (1^st^ vs 2^nd^ and 3^rd^ recording sessions; **Fig. 6**). We further conducted correlation analysis between LFP power (**Fig. 6**) and BOLD signal profiles (**Fig. 3** and **Supplementary Fig. 6**). Relationship between the differences of LFP power and BOLD signal profiles showing neural adaptation across multiple stimulation sessions were described by correlation coefficient, r and corresponding P value. The respective r and P values upon the three distinct stimulations were OB: r = 0.77 (P = 0.01), AON: r = 0.46 (P = 0.13), and Pir: r = - 0.06 (P = 0.85; **Supplementary Fig. 8**). This finding indicates that the decrease in LFP power (i.e., neural adaptation) is highly correlated with the decrease in BOLD activations, specifically upon OB or AON stimulation compared to Pir stimulation.

**Figure 5.**
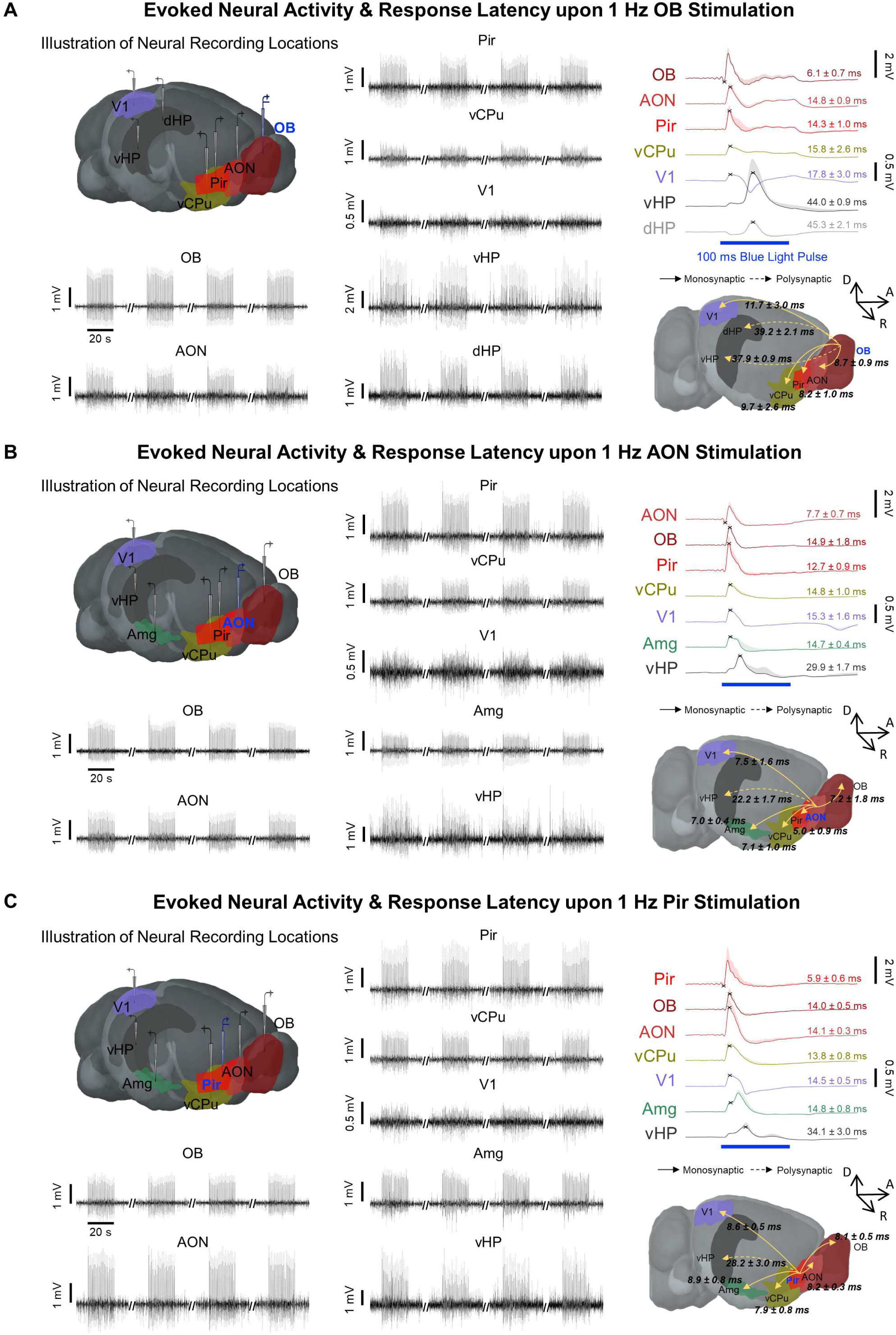
Local field potential (LFP) recordings confirm neuronal activity underlying BOLD activations and reveal orthodromic neural activity propagation to downstream targets in primary olfactory, striatal, visual, limbic and hippocampal networks. Illustration of electrophysiological recording regions (Top-Left), averaged evoked LFPs from the seven recorded regions (Bottom-Left & Middle), and summary of orthodromic neural activity propagation latencies measured from the stimulation onset over mono- and poly-synaptic projections (Right) upon 1 Hz optogenetic stimulation of (**A**) OB excitatory neurons, (**B**) OB afferents at AON and (**C**) OB afferents at Pir (n = 7, n = 7 and n = 5, respectively; 10% duty cycle; error bars indicate ± SEM; crosshairs denote LFP peaks that represent initial neuronal population activity for latency measurements). Significant delay between AON/Pir and OB neuronal population activity with appreciable jitter showed that neural activity propagation through the excitations of OB axonal terminals at AON or Pir, respectively, were largely orthodromic in our experiment.

**Figure 6.**
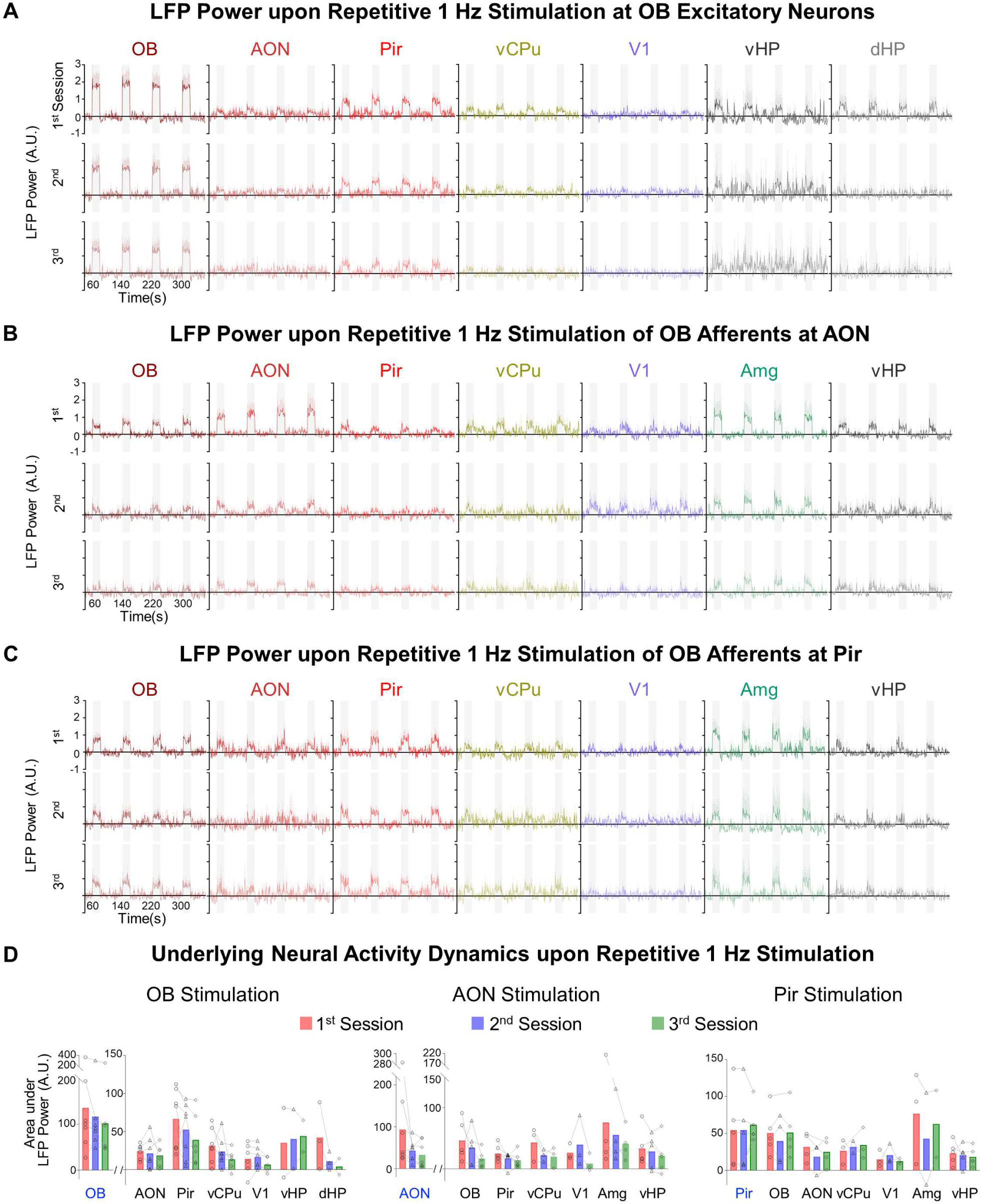
LFP recordings confirm varied neural activity adaptation in olfactory networks upon repetitive optogenetic excitations of OB excitatory neurons and OB afferents at AON and Pir. LFP power of evoked neural activities upon 1 Hz optogenetic stimulation of (**A**) OB excitatory neurons (n = 7; 10% duty cycle; error bars indicate ± SEM), (**B**) OB afferents at AON (n = 7; 10% duty cycle; error bars indicate ± SEM) and (**C**) OB afferents at Pir (n = 5; 10% duty cycle; error bars indicate ± SEM) across three electrophysiological recordings sessions. (**D**) Summary of the area under LFP power to quantify the extent of neural activity adaptation in the recorded regions of olfactory networks (individual animal data points are shown). In accord with our fMRI findings, the strength of neural activities, which were quantified through LFP power analysis here (i.e., area under absolute LFP trace normalized to 30-s baseline LFP power), was also decreased under repetitive 1 Hz stimulation of OB and OB afferents at AON but was relatively unchanged under Pir stimulation (1st vs 2nd and 3rd recording sessions). The LFP recordings demonstrate strong neural activity adaptation mediated by AON-driven neural activities to downstream targets.

We next characterized the latency of neural activity at each recorded region by using the first appreciable evoked LFP peak to represent the initial neuronal population activity^28,29,46^ (**Fig. 5**). Negative LFP peaks occurred first in each of the three stimulation targets, while positive peaks were observed for their corresponding downstream targets. During somatic activation of ipsilateral OB excitatory neurons, there is an evoked LFP response latency of 6.1 ± 0.7 ms (**Fig. 5A**), which is within the 5-6 ms range documented for response latency of OB excitatory neurons in the presence of natural odors^47,48^. We measured delays of 8.7 ± 0.9 ms and 8.2 ± 1.0 ms between OB and AON, and OB and Pir, respectively, consistent with the reported 5-10 ms neural activity propagation along monosynaptic projections of OB to AON^49^ and Pir^50–52^. Neural activities from ipsilateral OB arrived at ipsilateral vCPu and V1 in 9.7 ± 2.6 ms and 11.7 ± 3.0 ms in line with the axonal conduction speed between OB and striatal^53,54^/sensorimotor cortical^55,56^ regions. Lastly, the long delays of 37.9 ± 0.9 ms and 39.2 ± 2.1 ms from ipsilateral OB to vHP and dHP respectively, indicate that neural activities propagated through polysynaptic projections of OB-primary olfactory cortices-Ent-HP^52,57^. When stimulating the OB afferents in AON, we found a response latency of 7.7 ± 0.7 ms locally at AON (**Fig. 5B**). A subsequent delay of 7.2 ± 1.8 ms and 5.0 ± 0.9 ms was then measured between AON and its downstream targets in the primary olfactory network (i.e., OB and Pir), corroborating reported values of 4-10 ms^58^. The presence of a significant delay between AON and OB neuronal population activities with appreciable jitter demonstrates that neural activity propagation through the activation of OB axonal terminals at AON is largely orthodromic in our experiment. Neural activity from olfactory-specific AON neurons also propagated in an orthodromic manner to targets in the striatal (vCPu), sensorimotor cortical (V1) and limbic (Amg) networks, 7.1 ± 1.0 ms, 7.5 ± 1.6 ms, and 7.0 ± 0.4 ms, respectively, in line with axonal conduction speed^59^. Meanwhile, excitation of the OB afferents at Pir evoked a response latency of 5.9 ± 0.6 ms (**Fig. 5C**), which closely matched the latency of ∼5 ms upon optogenetic stimulation of afferents to Pir in a previous study^42^. Subsequently, the neural activity propagated from Pir to OB, AON, vCPu, V1, and Amg at a delay of 8.1 ± 0.5 ms, 8.2 ± 0.3 ms, 7.9 ± 0.8 ms, 8.6 ± 0.5 ms, and 8.9 ± 0.8 ms, respectively. We conclude that the observed neural activity propagation is predominantly orthodromic given the significant delay and jitter of excitatory inputs from Pir to OB. Notably, the neural activity propagated in 28.2 ± 3.0 ms from Pir to vHP, while the propagation speed from the AON to vHP was faster at 22.2 ± 1.7 ms. The longer delay between Pir and the hippocampus suggests the presence of an intermediary region between the Pir and Ent, such as the perirhinal cortex^60,61^.

Taken together, our electrophysiological recordings showed the direct involvement of both local and long-range projections that mediate the propagation of neural activity in olfactory networks. Moreover, orthodromic propagations when stimulating OB afferents at AON and Pir indicate that the long-range inhibitory effect of AON and excitatory effect of Pir outputs, which shaped the dynamic properties of BOLD activations in olfactory networks, are predominantly driven by the feedforward projections of olfactory-specific neurons in the primary olfactory cortices. Note that while we can also apply DCM analysis to our LFP recordings as we did for the fMRI data (**Fig. 4**) to examine the inhibitory effects of AON, we were unable to do so due to the lack of electrophysiological recording at Ent.

### Long-range olfactory networks of aged brains exhibit decreased neural activations and impaired AON to Pir connectivity

Informed by the insights gained from characterizing the spatiotemporal properties of long-range olfactory networks in normal healthy brains, we deployed our optogenetic fMRI platform to examine the systemic dysfunction of olfactory networks in a diseased brain. Here, we employed a D-galactose model of accelerated aging in rodents. This model induces higher levels of oxidative stress, mitochondrial dysfunction, and cellular apoptosis. These factors can lead to the decline of cognitive functions, which is a key symptomatic hallmark of age-related brain senescence^62,63^. Spatially, we observed that 1 Hz stimulation of the OB excitatory neurons evoked bilateral activations in all brain regions like primary olfactory, limbic, hippocampal, striatal and sensorimotor networks (**Fig. 7A** vs. **Fig. 2A**). These spatially preserved activations indicate that neural activity propagation in long-range olfactory networks remains intact in aged brains. However, the strength of neural activities at downstream activation targets exhibited a general decrease in both brain hemispheres when compared to healthy animals. A significant decline in BOLD activations was mostly identified in ipsilateral primary olfactory (aged vs. healthy; AONR: 16.4 ± 3.8 vs. 33.4 ± 6.5, *P* < 0.05; PirR: 33.5 ± 6.0 vs. 58.0 ± 8.4, *P* < 0.05; TTR: 15.1 ± 4.8 vs. 41.3 ± 7.0, *P* < 0.01) and ipsilateral limbic networks (CgR: 7.6 ± 1.9 vs. 15.5 ± 1.9, *P* < 0.01; OFCR: 19.4 ± 3.0 vs. 36.7 ± 3.8, *P* < 0.01; InsR: 29.2 ± 3.8 vs. 40.6 ± 5.0, *P* < 0.05). We also observed a general trend of decline in the other regions of the primary olfactory network (PirL: 23.7 ± 5.4 vs. 37.8 ± 7.1, *P* = 0.07 and EntR: 26.0 ± 4.8 vs. 38.3 ± 5.7, P = 0.06), and the ipsilateral sensorimotor network (MCR: 16.5 ± 2.4 vs. 24.5 ± 4.3, *P* = 0.059). Surprisingly, there was a general trend of increased BOLD activations in the striatal network (NAcR: 27.7 ± 2.8 vs. 20.3 ± 3.2, *P* = 0.055). DCM of the optogenetic fMRI data in aged animals revealed a switch in the effect of ipsilateral AON outputs to ipsilateral Pir from excitatory to inhibitory (healthy vs. aged; 0.23 ± 0.09 vs. −0.12 ± 0.10 Hz, *P* < 0.05), indicating dysfunction of a key primary olfactory cortical circuit in the olfactory network disrupting neural activity propagation to primary olfactory and limbic networks. Note, that in healthy animals (**Fig. 2D**), the primary olfactory network was preferentially recruited by AON-driven neural activities, while the limbic network was recruited mainly by Pir-driven neural activities. The predominant effect observed in the right hemisphere was likely due to the stimulation of the right/ipsilateral OB. We expect both hemispheres to be affected equally in aged brains.

**Figure 7.**
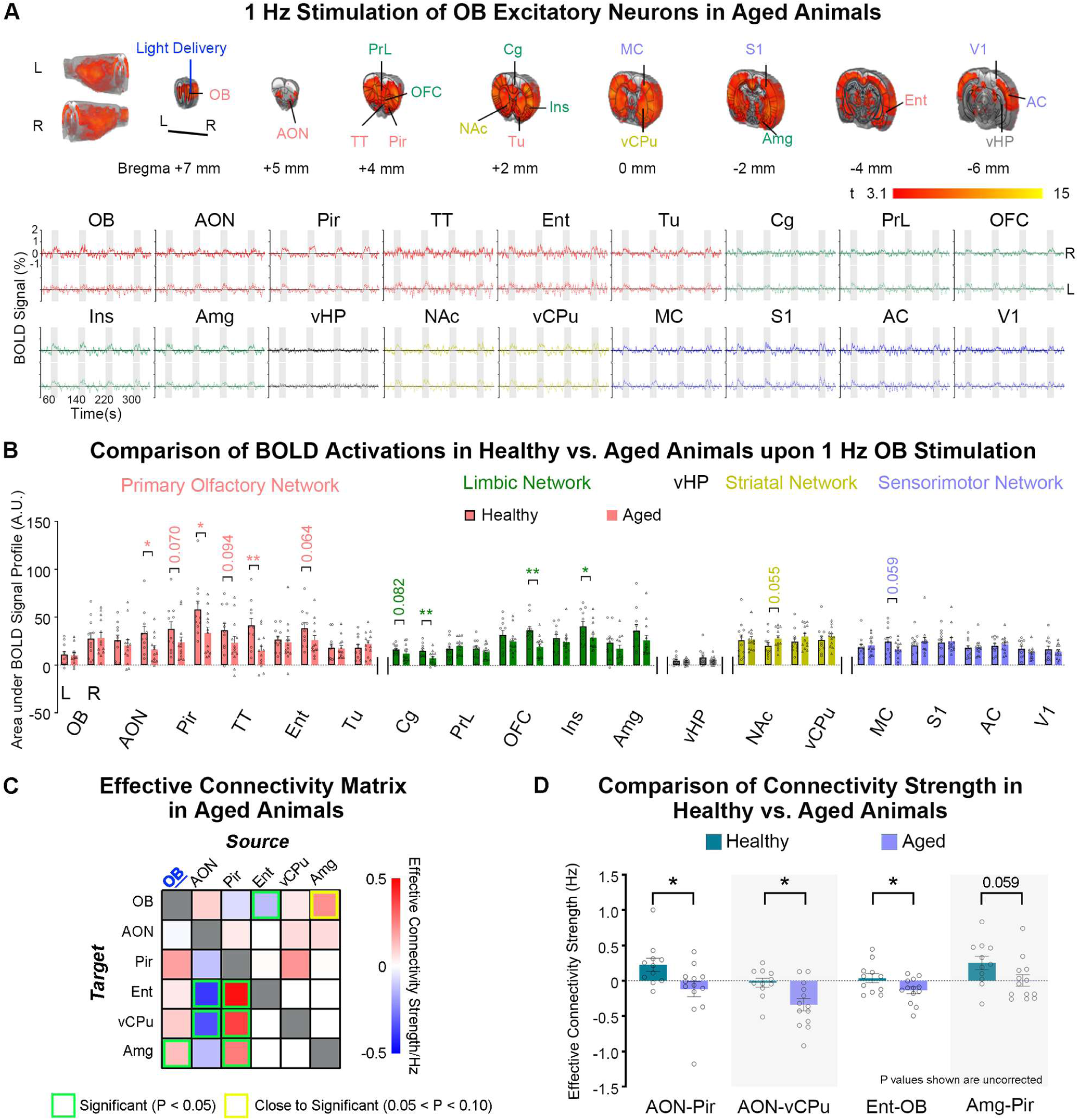
Decreased BOLD activations in primary olfactory and limbic networks, and impaired AON to Pir connectivity in aged animals. **(A)** Averaged BOLD activation maps and signal profiles from the 1^st^ fMRI session upon 1 Hz stimulation of OB excitatory neurons in D-galactose aged rat model (n = 13; t > 3.1 corresponding to *P* < 0.001; error bars indicate ± SEM). Note that the BOLD activation maps displayed in A were further corrected for multiple comparisons with TFCE-FWE at *P* < 0.05. (**B**) Comparison of the corresponding area under the BOLD signal profiles for each atlas-defined region-of-interests between healthy and aged animals (individual animal data points are shown; Fisher’s LSD test; \*\**P* < 0.01, and \**P* < 0.05; error bars indicate ± SEM). (**C**) Effective connectivity matrices of the olfactory network model upon stimulation of OB excitatory neurons in aged brains. (**D**) Quantitative summary of the significant differences in connectivity strength of downstream connections identified from the statistical comparison of effective connectivity matrices between healthy and aged brains (individual animal data points are shown; unpaired t-test; \**P* < 0.05; error bars indicate ± SEM). We uncovered a switch in the polarity from positive to negative in the connectivity of AON neural activity outputs to Pir in aged animals, indicating the dysfunction of a key primary olfactory cortical circuit that likely caused the significantly decreased BOLD activations in the primary olfactory and limbic networks.

## Discussion

As a major sensory system, the olfactory system has been linked to numerous non-olfactory regions pivotal for sensory perception and higher-order cognition^5^, yet recent views highlight the significant gap in understanding where and how olfactory information is distributed downstream beyond these known primary regions^5,64–67^. In this study, we deployed optogenetic stimulation, fMRI, and electrophysiological recordings to systematically dissect the spatiotemporal properties of long-range olfactory networks. We discovered that olfactory-specific neural populations in AON and Pir distinctly recruited downstream targets such as the hippocampal and striatal networks for AON-driven neural activities, and the limbic network for Pir-driven neural activities. Further analysis revealed robust adaptation of brain-wide BOLD fMRI activations upon repeated optogenetic stimulations of OB afferents in AON or OB excitatory neurons, which were absent when stimulating OB afferents at Pir. Computational modeling of our optogenetic fMRI data through DCM indicated a significant inhibitory effect of AON outputs to downstream targets in the olfactory network compared to the excitatory effect from Pir outputs. These results highlight a likely dominant role of olfactory-specific AON neural populations in mediating sensory adaptation in the presence of olfactory inputs. Electrophysiological recordings demonstrated predominant orthodromic neural activity propagation to downstream targets in long-range olfactory networks for all optogenetic stimulations and verified the adaptation of optogenetically-evoked neural activities observed in the fMRI experiments. In addition, our systematic comparisons of the olfactory networks between aged and healthy brains revealed an overall decrease in the amplitudes of BOLD activations, particularly in regions of the primary olfactory and limbic networks, along with increased inhibitory effect of AON outputs to the Pir, indicating impairment of this key olfactory cortical circuit.

While we cannot discount that certain characteristics of brain states differ under anesthetized and awake conditions, we showed that light (1.0% isoflurane) anesthesia minimally affects the BOLD fMRI activations and/or the propagation of optogenetically-evoked neural activity across thalamo-cortical, hippocampal-cortical and vestibulo-cortical networks^28–31,46,68,69^. Thus, our optogenetic fMRI approach constitutes the initial characterization of 1) the spatiotemporal properties of long-range olfactory networks and 2) the roles of AON and Pir outputs in shaping the dynamic properties of olfactory neural activity propagation in healthy and diseased brains upon optogenetic stimulation of OB, AON and Pir. Note that we did not utilize natural odor stimulation in this study.

Habituation or adaptation shows decreased responses to repeated or continuous stimuli^70,71^. Olfactory adaptation refers to the ability of the network to adjust its sensitivity at different stimulus intensities, which is essential for retaining high sensitivity to odd/another stimulus under repetitive or prolonged odor stimulation^72^. This fundamental property of all sensory systems prevents overstimulation and retains alertness to the environment^71^. Olfactory adaption occurs at two levels: peripheral – at the level of olfactory receptors – and central – at the level of the central nervous system^71^. At present, the underlying neural mechanism of peripheral adaptation that occurs at the molecular level^72^, and central adaptation within the primary olfactory local network (i.e., OB^73,74^, AON^25^, Pir^19,75^ and Tu^76^) has been shown to require glutamate release in the OB^77^. However, the significant decreases in BOLD activation across all downstream targets indicate that olfactory adaptation may occur more broadly across various targets of the olfactory network in addition to being localized at the OB. Specifically, our results reveal a previously undefined role of the AON in mediating olfactory adaptation at the systems level, as we showed strong adaptation of optogenetically-evoked neural activities (**Figs. 3B** and **6B**) upon 1 Hz stimulation of OB projection neurons and OB afferent inputs at AON, but not OB afferents at Pir, along with robust inhibitory effects of AON outputs to Ent, vCPu and Amg (**Fig. 4C**).

AON sends robust centrifugal inputs to OB that can inhibit bulbar outputs^25,58^ and robust glutamatergic feedforward projections that bilaterally innervate the Pir^42,78^, to enhance activation of piriform neurons in response to odor^42^. Further, recent studies on AON functions revealed its involvement in interhemispheric communication^25,79,80^, sensory gating^81,82^ and odor memory^33,36^, indicating a predominant role of AON as a hub to process olfactory information (i.e., odor identity decoding and gain control of OB outputs^43^). In our study, only stimulation of the ipsilateral AON evoked bilateral brain-wide BOLD responses in primary olfactory (i.e., Pir, Tu), limbic (i.e., Ins, OFC) and striatal regions (i.e., NAc, vCPu) at all frequencies (**Fig. 2B and Supplementary Fig. 4**), indicating the pivotal role of AON in olfactory processing between both hemispheres. AON receives centrifugal projections from HP, which represent its ability to process odor memory-related contextual information^33,36,81,83,84^. Notably, our fMRI analysis also showed that AON-driven neural activities preferentially recruited the hippocampal network. Further, we observed robust bilateral brain-wide activations initiated by 1 Hz stimulation at AON during the first fMRI session, which were followed by dramatically decreased downstream activations under repetitive stimulations (**Fig. 3**). We speculate that AON exerts robust feedforward neural activity propagation to long-range downstream targets after the early stage of olfactory processing at OB. However, in the presence of repeated strong activations of olfactory-specific neural populations in AON (i.e., through OB or AON stimulations), centrifugal feedback from the AON, which preferentially controls the gain of OB neuronal responses^43^, likely inhibited OB outputs^25,58^, resulting in decreased brain-wide activations in later fMRI sessions. Further, we identified robust inhibitory AON outputs to non-primary olfactory brain regions (i.e., Ent, vCPu, Amg; **Fig. 4C**), regardless of the stimulated region, indicating that neural adaptation processes could also be distributed beyond local olfactory micro-circuits (i.e., centrifugal AON outputs to OB). We note that olfactory adaptation at the systems level is quite under-studied, and future studies may examine the circuit properties at these long-range targets when processing odor information.

Pir sends ipsilateral inputs to OB and AON^40,85^ and functions to amplify the inhibition of odor-related signals and finely tune them in OB^43,86^, in order to process and identify critical features of an odor. Although, studies have noted a significant overlap in the functions of Pir during olfactory processing with those of AON^5,38,42^, the recurrent neural architecture of Pir^87,88^ stands in contrast to that of AON. These structural differences indicate how Pir processes olfactory inputs from OB distinctly. We found that Pir-driven neural activities preferentially recruited regions in the limbic network, which differed from those primarily recruited by AON. Anatomically, Pir projects extensively to regions of the limbic network, such as OFC and Amg^89,90^, which demonstrates the specificity in our optogenetic fMRI approach when dissecting the spatiotemporal properties of long-range olfactory networks. Overall, we also observed that exciting olfactory-specific neurons in Pir produced weaker brain-wide BOLD activations than OB or AON stimulation (**Fig. 2C** and **Supplementary Fig. 5**). We posit this occurs from the recurrent neural architecture of Pir (i.e., dense reciprocal synaptic connections between olfactory-specific neurons in Pir and inhibitory interneurons), which can recruit sustained inhibition that subsequently diminishes overall Pir cortical neural activity in the presence of olfactory inputs^87^.

The ability to detect and discriminate odors declines during aging, leading to a decrease in quality of life and diminished appetite with a concomitant impact on nutrition^91^. Decreased olfactory function with aging parallels generalized age-related deficits in sensory functions and cognition that occur in the absence of obvious disease states^14^. However, studies in rodents demonstrate that the decreased ability to perceive odors was not associated with a general decline in the number of receptors/cells in the olfactory epithelium or a decline in neuronal populations (i.e., mitral and tufted cells, and interneurons) in the OB^13^. Instead, the synaptic density in the glomeruli of the OB appears diminished^13^, suggesting impaired downstream olfactory information processing from the OB to targets in the cortical and subcortical regions. At present, there is no consensus on the most representative animal model of aging that best mimics the human condition. D-galactose has been shown to induce advanced glycation end products formation, a process that contributes to early phases of age-related diseases in both humans and rodents^92,93^. With the discovery of diminished brain-wide neural activity propagation, our results in a D-galactose rat model indicate that the impairment of OB downstream olfactory information processing is likely true in aged brains. We also found the switch from positive effective connectivity of AON to Pir and AON to vCPu in healthy animals to significantly negative ones in aged animals, indicating abnormal AON output to downstream targets in long-range olfactory networks during olfactory information processing. Notably, age-related neurodegenerative diseases (e.g., Alzheimer’s and Parkinson’s disease) can exhibit olfactory impairments in the early stages with cell loss and axonal degradation in AON^94–96^. While the comparisons made here were not with age-matched healthy controls, they provide preliminary insights into aging-related dysfunctions of brain-wide olfactory networks to guide future studies.

In summary, our study demonstrates the distinct recruitment of higher-order brain networks by primary olfactory cortices (AON and Pir), including the identification of AON and Pir outputs that can shape the dynamic properties of neural activity propagation in long-range olfactory networks in healthy and aged brains. Such documentation of long-range olfactory networks and their properties will be of great value to both neuroscientists and clinicians. By gaining an integrated understanding of olfactory networks and their fundamental properties in both healthy and aged brains, we can develop therapeutic strategies for neurological diseases associated with olfactory system dysfunctions.

## Methods

### Animal subjects

Adult male Sprague Dawley rats were used in all experiments. Animals were individually housed under a 12-h light/dark cycle with access to food and water ad libitum. All animal experiments were approved by the Committee on the Use of Live Animals in Teaching and Research (CULATR) of the University of Hong Kong. Seven groups of animals were used: Group I ( n = 11) for optogenetic fMRI experiments upon olfactory bulb (OB) stimulation; Group II (n = 11) for optogenetic fMRI experiments upon anterior olfactory nucleus (AON) stimulation; Group III (n = 9) for optogenetic fMRI experiments upon piriform cortex (Pir) stimulation; Group IV (n = 7) for electrophysiological recordings upon OB optogenetic stimulation; Group V (n = 7) for electrophysiological recordings upon AON optogenetic stimulation; and Group VI (n = 5) for electrophysiological recordings upon Pir optogenetic stimulation; and Group VII (n = 13) for optogenetic fMRI experiments upon OB stimulation in aged rats.

Note that for the electrophysiological experiments, not all animals resulted in successful recordings in all electrode locations. For Group IV, only OB, AON, Pir, vCPu and V1 were recorded in all 7 animals, whereas only 2/7 in vHP and 3/7 animals in dHP. For Group V, AON in 7 animals, OB in 6 animals, whereas only 5/7 in Pir, 3/7 in vCPu, V1 and vHP, and 4/7 animals in Amg. For Group VI, OB, Pir and vHP in 5 animals, followed by 4/5 in AON, 3/5 in vCPu and V1, and 2/5 animals in Amg.

### Virus packaging

Recombinant AAV vectors were serotyped with AAV5-coated proteins and produced by the Vector Core of the University of North Carolina. Viral titer in particles/mL was 4 × 10^12^ for AAV5CaMKII *α*-ChR2(H134R)-mCherry. Maps are available online from http://www.stanford.edu/group/dlab/optogenetics.

### Accelerated aging animal model induction protocol

D-galactose (50 mg/kg, subcutaneous) was administered once daily to 10-week-old male adult SD rats for a total duration of 8 weeks. D-galactose administration in rodents has been shown to cause increased oxidative stress, mitochondrial dysfunction and cellular apoptosis, which leads to the deterioration of cognitive functions, a primary symptomatic hallmark of senescence of the brain^62,63^.

### Stereotactic surgery for viral injection

Stereotactic surgeries were performed with rats at 6-7 weeks of age for group I to VI, and at 15 weeks (i.e., 5 weeks in on the daily D-galactose administration) for group VII. Rats were anesthetized with an intraperitoneal bolus injection of a ketamine (90 mg/kg) and xylazine (40 mg/kg) mixture. After shaving the scalp, rats were secured in a stereotactic apparatus with non-rupturing ear bars and craniotomy was conducted. Buprenorphine (0.05 mg/kg, subcutaneous) was administered to minimize pain. A heating pad was used to prevent hypothermia. Viral injection was performed at two depths in OB (+7.5 mm anterior to Bregma, +1.8 mm medial-lateral right hemisphere, −1.8 mm and −2.1 mm from the surface of dura), with subsequent verification by high-resolution anatomic MRI. Viral constructs were delivered at 1.5 μL volume at each depth through a 5 μL syringe and 33-gauge beveled needle injected at a rate of 150 nL/min. The injection needle was held in place for 10 min and retracted from the brain slowly. Scalp incision was sutured, and animals were kept on a heating pad until recovery from anesthesia. Buprenorphine (0.05 mg/kg, subcutaneous) was administered post-injection twice daily for 72 hours to minimize discomfort. Enrofloxacin was also administered orally for 72 hours to minimize infection and inflammation post-surgery. Animals recovered for 4-5 weeks before conducting MRI and electrophysiology experiments.

### Animal preparation for optogenetic fMRI experiments

Before surgery, animals were anesthetized with isoflurane (induction 3% and maintenance 2%) and secured on a stereotactic frame. Before animals were placed in-magnet, surgery was performed under 1.2-2.0% isoflurane to implant a custom-made plastic optical fiber cannula (core diameter = 450 μm, outer diameter = 500 μm). Implantation of the cannula was made at three different locations in the ipsilateral/right brain hemisphere depending on the experiments, namely at OB (+7.5 mm anterior to Bregma, +1.8 mm medial-lateral, −2.1 mm from the surface of dura), AON (+5.2 mm anterior to Bregma, +1.8 mm medial-lateral, −4.3 mm from the surface of dura) and Pir (+4.2 mm anterior to Bregma, +3.1 mm medial-lateral right hemisphere, −5.4 mm from the surface of dura). Following a midline incision, a craniotomy was performed at these located coordinates. The fiber tip was scored to create a beveled edge to facilitate insertion of the fiber and minimize injury to brain tissue. The fiber cannula was then made opaque using heat shrinkable sleeves to prevent light leakage during stimulation and the eyes of the animals were blindfolded to prevent undesired visual stimulation, as in our previous studies^28–31,46,68^. Dental cement was applied to the fiber cannula to affix it to the skull. The optical fiber tip was typically situated in the mitral cell layer and external plexiform layer in OB stimulation experiments, dorsal pars principalis of the AON to target OB afferents at AON, and anterior Pir for OB afferents at Pir with verification by high-resolution anatomical MRI (**Fig. 1A**). The spatial spread from the fiber tip for 473 nm blue light is rather small (200 μm and 350 μm at 50% and 10% of initial light intensity, respectively) and has little or no in the backward and lateral directions^28,97^, confining the optogenetic stimulation to the excitatory neurons in OB, and the axonal terminals of OB at AON and Pir, respectively. Buprenorphine (0.05 mg/kg, subcutaneous) was administered post-implantation.

Following the surgical procedure for fiber cannula implantation, animal was immediately prepped for fMRI experiments. One drop of 2% lidocaine was applied to the chords to minimize discomfort before endotracheal intubation. The animals were mechanically ventilated at a rate of 60 breaths per minute with 1.0% isoflurane in room-temperature air using a ventilator (TOPO, Kent Scientific, Torrington, CT). During all fMRI experiments, animals were placed on a plastic holder, and their heads were fixed with a tooth bar and plastic ear bars with screws. Continuous physiological monitoring was performed using sensors from an MRI-compatible system (SA Instruments, Stony Brook, NY). Rectal temperature was maintained at ∼37.0 °C using a water circulation system. Vital signs were within normal physiological ranges (rectal temperature: 36.5-37.5 °C, heart rate: 350-420 beats/min, breathing: ∼60 breaths/min but not synchronized to optogenetic stimulation, oxygen saturation level: >95%) throughout the experiments.

### MRI scanner-synchronized optogenetic stimulation

An Arduino programming board synchronized the scanner trigger and the lasers for optogenetic stimulation. Computers and light delivery systems were kept outside the magnet, and optical patch cables (5-10 m) delivered light into the bore of the scanner. For optogenetic stimulation, blue light was delivered using a 473 nm Diode-pumped solid-state laser, measured before scanning as 8 mW at the fiber-tip (450 μm, NA = 0.5) corresponding to a light intensity of 40 mW/mm^2^. For all fMRI experiments (i.e., OB, AON or Pir stimulation), five frequencies were used (1 Hz with a 10% duty cycle; 5, 10, 20 and 40 Hz with a 30% duty cycle). OB, AON or Pir was stimulated with a block design paradigm that consisted of four 20s-on and 60s-off periods with the order of stimulation frequencies pseudorandomized (**Fig. 1B**). Typically, three fMRI sessions were performed for each animal (i.e., five scans per session covering the five different frequencies). The intervals between two fMRI scans were within one minute. Since the stimulation frequencies (i.e., 1 Hz, 5 Hz, 10 Hz, 20 Hz and 40 Hz) were pseudorandomized, the interval between any two identical stimulation frequency was on average 30 minutes apart.

### MRI acquisition procedure

All MRI experiments were carried out on a 7T MRI scanner (ParaVision v5.1, PharmaScan 70/16, Bruker Biospin GmbH, Ettlingen, Germany) using a transmit-only birdcage coil in combination with an actively decoupled single-channel receive-only surface coil. Scout T2-weighted (T2W) images were first obtained using a rapid acquisition with relaxation enhancement (RARE) sequence for accurate positioning and as an anatomical reference with field of view (FOV) = 32 × 32 mm^2^, matrix = 256 × 256, RARE factor = 8, echo time (TE) = 36 ms, and repetition time (TR) = 4200 ms. Seventeen contiguous 1 mm slices were then positioned in the transverse orientation according to the rat brain atlas to cover most of the brain from the OB to the cerebellum. All fMRI measurements were made using a single-shot gradient-echo echo-planar imaging (GE-EPI) sequence with FOV = 32 × 32 mm^2^, matrix = 64 × 64, flip angle = 56°, and TE/TR = 20/1000 ms.

### fMRI data analysis

fMRI analysis was performed using the procedure established in our previous studies^28–31,46,68^. For each fMRI session, all GE-EPI images were first corrected for slice timing differences and then realigned to the mean image of the first fMRI scan using the SPM12 fMRI toolbox (Wellcome Department of Imaging Neuroscience, University College London, UK). fMRI scans suffering from motion artifacts (i.e., voxel shifts > 0.05 mm detected by realignment) and sudden physiological changes (i.e., abrupt changes in respiration pattern, heart rate, and oxygen saturation level) were discarded. After resampling EPI and T2 images to 128 × 128 × 17, the EPI images from each animal were first registered to their respective T2W images. T2W images from each animal were then co-registered to a custom-made brain template acquired from separate age-matched rats with identical T2 scanning parameters using affine transformation and Gaussian smoothing to maximize normalized mutual information (SPM12). The transformation matrix was then applied to co-register the fMRI EPI volumes. Voxel-wise linear detrending with least-squares estimation was subsequently performed temporally for the EPI volumes to eliminate any baseline drift caused by physiological noise and/or system instability. For individual animals, data from the three fMRI sessions were smoothed (full width at half maximum, FWHM = 1 pixel with no out-of-plane smoothing), and high pass filtered (128 s). Then, a general linear model (GLM) was applied to calculate the response coefficient (β) maps for each stimulus. The optogenetic stimulation block-designed paradigm (**Fig. 1B**) were convolved with the canonical hemodynamic response function (HRF) and subsequently used as a regressor in GLM. Student’s t-test was performed to identify activated voxels using the threshold *P* < 0.001 (t value > 3.1). BOLD signal profiles were then extracted from regions of interest (ROIs) delineated from the rat brain atlas for further evaluation. These steps performed at the individual animal level were done to ensure that visual inspection and comparison of the data quality across animals could be conducted before group-level analysis. At the group level, resliced, realigned and co-registered EPI images corresponding to the same fMRI session and stimulation frequency were averaged across animals as follows. BOLD activation map of individual animal was first generated. These individual activation maps then underwent a two-step statistical thresholding method to generate the group-level results. Firstly, uncorrected one-sample t-tests were conducted with a threshold of P < 0.001. Secondly, these thresholded activation maps were further corrected using nonparametric inference with threshold-free cluster enhancement multiple comparison correction of family-wise error rate (TFCE-FWE, P < 0.05) before they were averaged. For the third fMRI session in experiments stimulating the AON and Pir, TFCE-FWE was not applied in the masking of thresholded BOLD activation maps were only masked with the group-level masks generated from one-sample group-level t-tests without TFCE-FWE corrections (no activations after t-tests and TFCE-FWE corrections, t > 3.1). BOLD signal profiles for each ROI were generated by averaging those extracted earlier from individual animals.

In the fMRI experiments upon 1 Hz stimulations of OB, AON and Pir, quantification of the strength of neural activity at downstream activation targets was performed separately for the three consecutive fMRI sessions in each animal. First, the cumulative area under the BOLD signal profile (AUC) for each atlas-defined ROI was computed. Here, only the area during the four 20-s-on period of optogenetic stimulation was considered. Given that the BOLD response onset in our experiments was ∼3 s after the presentation of the stimulus, we systematically delayed our area under BOLD signal profile measurements by 3 s. Besides, AUC offers a more comprehensive measure of BOLD signal change over time, including shape, duration, and magnitude, thereby capturing the bulk of neural activities and their dynamics throughout the stimulation period. Meanwhile, β primarily represents the peak amplitude of BOLD responses (i.e., the % BOLD signal change)^98^ and can be constrained by the assumptions and limitations of the GLM analysis, such as the shape of the canonical hemodynamic response function (HRF). Second, the computed AUCs were grouped into five networks such as the primary olfactory (OB, AON, Pir, tenia tecta, TT, entorhinal cortex, Ent and olfactory tubercle, Tu), limbic (cingulate, Cg, orbitofrontal cortex, OFC, insular, Ins and amygdala, Amg), striatal (nucleus accumbens, NAc and ventral caudate putamen, vCPu), hippocampal (ventral hippocampus, vHP) and sensorimotor (motor, MC, somatosensory, S1, auditory, AC and visual, V1, cortices). Third, BOLD activation strength for each defined network was then measured. Here, the AUC of each network was normalized to the AUC of all five networks. Lastly, the BOLD activation strengths for each network were statistically compared between the three different fMRI stimulation experiments targeting OB excitatory neurons and OB afferents at AON and Pir, respectively (outlier detection with nonlinear regression & false discovery rate at 1 %, Tukey’s multiple comparisons test). Note that direct comparisons between AUCs of each downstream target from the three separate optogenetic fMRI experiments were not considered, as they are likely erroneous given the differences in absolute BOLD signal amplitudes. For example, BOLD signal amplitudes in Pir stimulation experiments were generally weaker than OB and AON stimulations. The variations here are likely caused by differences in ChR2 transfection volumes in OB excitatory neurons and OB afferents at AON/Pir, including the population size of neurons/axonal terminals stimulated. Note that the vast majority of OB neurons are interneurons, and not projection neurons to OB and AON^38^.

Comparison of BOLD power between fMRI sessions (i.e., 1^st^ vs. 2^nd^, 1^st^ vs. 3^rd^ fMRI session) upon 1 Hz stimulation was done using Dunnett’s multiple comparisons test to evaluate olfactory adaptation under repetitive OB/AON/Pir stimulation. Normalized BOLD power was only shown for display (not all data points were shown, because some values were extremely huge) with statistical results calculated based on true BOLD power comparison (**Supplementary Fig. 6**). When comparing the BOLD power between healthy vs. aged rats upon first 1 Hz OB stimulation, we conducted an uncorrected Fisher’s LSD test (**Fig. 7B**).

### Spectral dynamic causal modeling (sDCM), and statistical analyses

To parametrize the downstream effects of OB-, AON- and Pir-driven neural activities on selected targets in the right brain hemisphere (i.e., same hemispheric side to the optogenetically-stimulated region) in healthy and aged animals, we further analyzed the BOLD fMRI data upon 1 Hz stimulation using sDCM^26^. A six-node brain network model was defined, encompassing connections between regions in the primary olfactory (OB, AON and Pir), striatal (vCPu) and limbic (Amg) networks (**Fig. 4A**). Specifically, sDCM was utilized to computationally model the effective connectivity strength of these downstream connections in OB, AON and Pir stimulation of healthy animals, and OB stimulation data of aged animals. To avoid over-fitting of the data, the BOLD signals were band-pass filtered with a low cutoff frequency at 1/128 Hz and a range of high cutoff frequencies at 0.1-0.3 Hz. We then evaluated the connectivity parameters from the choice of 21 high cutoff frequencies with a step size of 0.01 Hz. We selected the optimal high cutoff frequency for each stimulation target (i.e., OB, AON, Pir) that maximized the weighted sum of scores for all modeled connections (significant, *P* < 0.05, score = 1; close to significant, 0.05 < P < 0.1, score = 0.5; FDR corrected). The optimal high cutoff frequencies were 0.1, 0.18 and 0.14 Hz for OB, AON and Pir stimulations, respectively, in healthy animals, and 0.1 Hz for OB stimulation in aged animals. Stochastic sDCM has been demonstrated to perform better compared to deterministic modeling^26^. Here, we utilized Bayesian model selection to determine the autoregressive model order (e.g., autoregressive models of order one, AR1; order two, AR2), which was used to parameterize neural fluctuations and observed noise in stochastic models. We then applied Bayesian model averaging (BMA) to generate the mean effective connectivity for each defined connection since we did not obtain a clear winning autoregressive model order. Following BMA, the mean effective connectivity was statistically compared across animals and summarized in the form of connectivity matrices (**Fig. 4B**, n = 11, n = 11 and n = 9 for OB, AON and Pir stimulation in healthy animals; n = 13 for OB stimulation in aged animals; one-sample t-test with FDR correction). We further quantified the differences in the effective connectivity of AON-related vs. Pir-related connections to downstream targets such as Ent, vCPu and Amg by conducting paired t-test with FDR correction (**Fig. 4C**). Unpaired t-test was used to compare the effective connectivity of AON-related and Pir-related connections between healthy and aged animals (**Fig. 7D**).

### Multi-site local field potential (LFP) recordings and analyses

Electrophysiological recordings were performed under the same anesthesia protocols and similar physiological conditions as in the fMRI experiments with body temperature maintained using a heating pad. Guided by the fMRI findings, craniotomies were conducted and dura matter was removed with reference to the ROIs of ipsilateral OB (+7.5 mm anterior to Bregma, +1.8 mm ML), AON (+5.2 mm anterior to Bregma, +1.8 mm ML), Pir (+4.2 mm anterior to Bregma, +3.1 mm ML), vCPu (+2 mm anterior to Bregma, +2.7 mm ML), Amg (−3 mm posterior to Bregma, +3.8 mm ML)/ V1 (−6 mm posterior to Bregma, +4.2 mm ML), vHP (−6.4 mm posterior to Bregma, +6.4 mm ML) / dHP (−3.6 mm posterior to Bregma, +2.4 mm ML). For coverage of the majority of layers in the dentate gyrus of ipsilateral vHP/dHP, a linear microelectrode silicon array (16 recording channels equally spaced at 100 μm; 1.5 MΩ impedance; NeuroNexus Technologies, Ann Arbor, MI) was inserted perpendicularly to the brain surface with the tip of the linear array approximately 4.5 mm/3.7 mm below the dura matter via micromanipulators (Narishige, Amityville, NY). For other regions, single tungsten microelectrodes with 2.0 MΩ and 10 μm tip diameter were utilized (OB: −1.8 mm DV; AON: −4.3 mm DV; Pir: −5.4 mm DV; vCPu: - 5.4 mm DV; V1: −1.1 mm DV; Amg: −8.4 mm DV). Fiber and a single-channel electrode were constructed into an optrode^99^ and were implanted following identical procedures as described for optogenetic fMRI experiments. LFP data were acquired using a multi-channel neurophysiology recording system (sampling frequency: 24 kHz; notch-filter: 50 Hz, 100 Hz, and 150 Hz; Tucker Davis Technologies/TDT, Alachua, FL). Synchronized laser stimulation was controlled by the same system, and the trigger signal for blue light pulse trains was recorded simultaneously with the extracellular recording data. Experimental paradigms were identical to those used in fMRI experiments.

After recordings, raw data were bandpass-filtered (0.01-150 Hz) and down-sampled to 1 kHz using MATLAB. Individual LFP and group-averaged LFP traces were generated. Averaged LFP traces were presented with standard error mean to reflect the reproducibility across animals as shown by our previous analyses and literature^28,29,46,100–102^. The latency of evoked LFPs was measured from the onset of light pulse to the first appreciable peak location (i.e., either negative or positive). To quantify the optogenetically evoked neural activity power during the 20-s optogenetic stimulation block, the area under absolute LFP traces was measured as LFP power during the optogenetic stimulation. Given that the LFP response onset was a few milliseconds after the presentation of the stimulus, no time delay for the area under the LFP power measurements (**Fig. 6**).

Correlation coefficient analysis between LFP power and BOLD signal profiles were also conducted. First, for each 1 Hz optogenetic stimulation target (i.e., OB, AON, and Pir), ROIs that have both LFP recordings and BOLD activations were selected. The selected ROIs were AON, Pir, vCPu, V1, vHP upon OB stimulation; OB, Pir, vCPu, V1, Amg, vHP upon AON stimulation; and OB, AON, vCPu, V1, Amg, vHP upon Pir stimulation. Note that the respective stimulated region was excluded due to BOLD fMRI signal dropout caused by the implanted optical fiber. Second, the AUC differences of 2nd stimulation session vs. 1st session and 3rd session vs. 1st session in LFP power and BOLD signal profiles of individual animal were calculated, respectively. Subsequently, the mean AUC differences for each ROI across animals were calculated. Third, the correlation coefficients and P values of AUC differences between LFP power and BOLD signal profiles were calculated using Spearman nonparametric correlation (**Supplementary Fig. 8**).

### Histology, immunohistochemistry, and confocal imaging

To confirm the specific expression of ChR2-mCherry in the OB excitatory neurons and the axonal inputs to AON/Pir, histology was performed according to the previously published procedure^28–31,46,68^. Upon completion of experiments, a selected number of optogenetically transfected animals (n = 3 for OB; n = 2 for AON/Pir) were anesthetized with pentobarbital and then transcardially perfused with ice-cold 4% paraformaldehyde (PFA) in phosphate-buffered saline (PBS). The brains were equilibrated in 20% sucrose in PBS at 4 °C overnight. Axial sections (40 μm) were prepared on a freezing microtome (model 860, AO Scientific Instruments). Consecutive sections were mounted and examined with a laser confocal microscope (Carl Zeiss LSM780/LSM980). For immunohistochemistry, free-floating sections were processed with 5% normal goat serum and 0.3% Triton X-100 in PBS with primary antibodies against rabbit polyclonal to CaMKIIα (1:400; Abcam) at 4 °C for 24 h. After washing with PBS, sections were then incubated for 2 h at room temperature with secondary antibodies Alexa Fluor 647 conjugate goat anti-rabbit IgG and Alexa Fluor 488 conjugate goat anti-guinea pig IgG (both 1:500; Molecular Probe). Slices were then washed and mounted using FluoroShield mounting medium with DAPI (Abcam). Double or triple immunofluorescence was assessed with a laser confocal microscope (Carl Zeiss LSM780/LSM980).

## Data availability

The fMRI and electrophysiology data generated in in this study are under active use by the reporting laboratory; all the raw data that support the findings of this study are available from the corresponding author upon request. The respective data for line graphs, bar charts, and matrices in the main figures (Figs. 2-7) are provided with this paper in the **Source Data 1** file.

## Code availability

All custom MATLAB software codes used for fMRI and LFP analyses, and DCM analysis in this study can be downloaded from the **Source Code 1** file provided. Other information is available from the corresponding author upon reasonable request.

## Acknowledgement

This work was supported by the Hong Kong Research Grant Council (HKU17130225, HKU17127723 and HKU17127021 to A.T.L. Leong, and C7052-23GF, HKU17130325, HKU17119224, HKU17127523 and HKU17127022 to E.X. Wu) and Lam Woo Foundation to E.X. Wu. We would like to thank Drs. L.W. Lim, P.S. Chong, E. Wong, and Ms. S. Kwok for their advice and/or technical assistance. We would also like to thank Dr. K. Deisseroth who provided us with the ChR2 viral construct.

## Author Contributions

Study design and conceptualization: T.M., X.W., A.T.L.L. and E.X.W.. Animal fMRI experiments: T.M.. Animal fMRI data analysis: T.M., X.W., A.T.L.L. and X.L.. Electrophysiology experiments: T.M. and J.W.. Electrophysiology data analysis: T.M., X.W. and A.T.L.L.. Visualization: T.M., A.T.L.L., X.W., X.L. and L.X.. Suggestions: X.W., A.T.L.L. and L.X.. Supervision: A.T.L.L., E.X.W., X.W. and P.C.. Original draft writing: T.M. and A.T.L.L.. Writing-review and editing: A.T.L.L., E.X.W., T.M. and X.W..

## Additional Information

Figure supplements accompany this paper.

## Supplemental Information (SI)

**Supplementary Figure 1.**
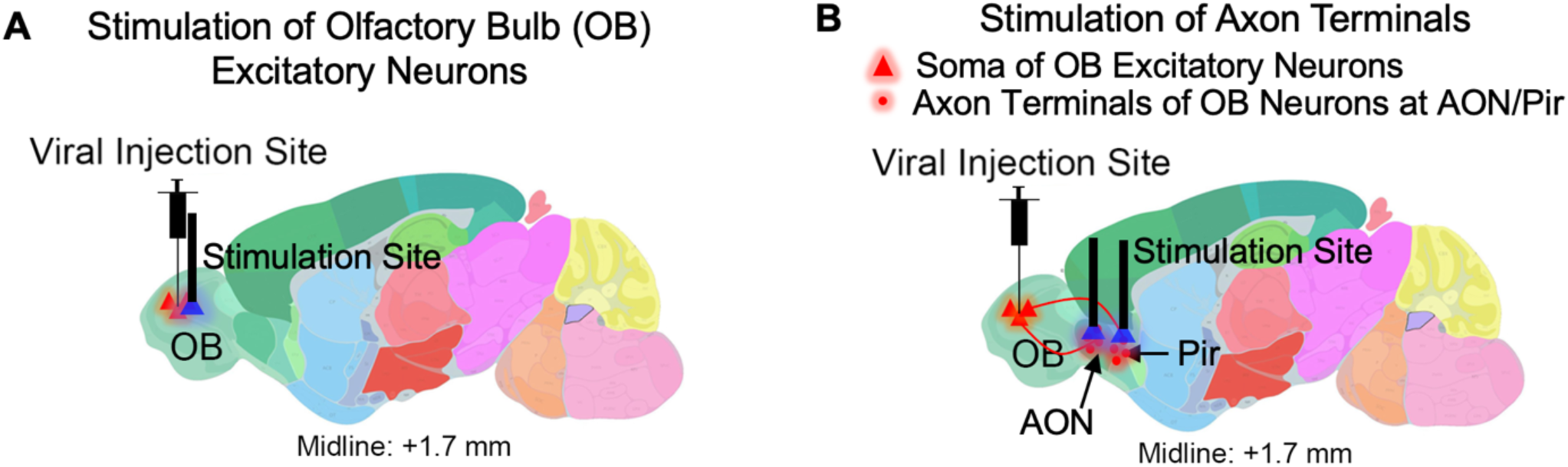
Illustration of the optogenetic stimulation protocols implemented in the study. (**A**) Optogenetic stimulation of the cell bodies/soma of CaMKIIα::ChR2 expressing excitatory olfactory bulb (OB) neurons. (**B**) Optogenetic stimulation based on projection targeting technique. Here, the ChR2 expressing axon terminals of excitatory OB neurons (i.e., OB afferents) at primary olfactory cortices, such as anterior olfactory nucleus (AON) and piriform cortex (Pir), were stimulated instead. Note that the three different stimulations (i.e., OB excitatory neurons, OB afferents at AON and OB afferents at Pir) illustrated here were conducted separately in different animal groups.

**Supplementary Figure 2.**
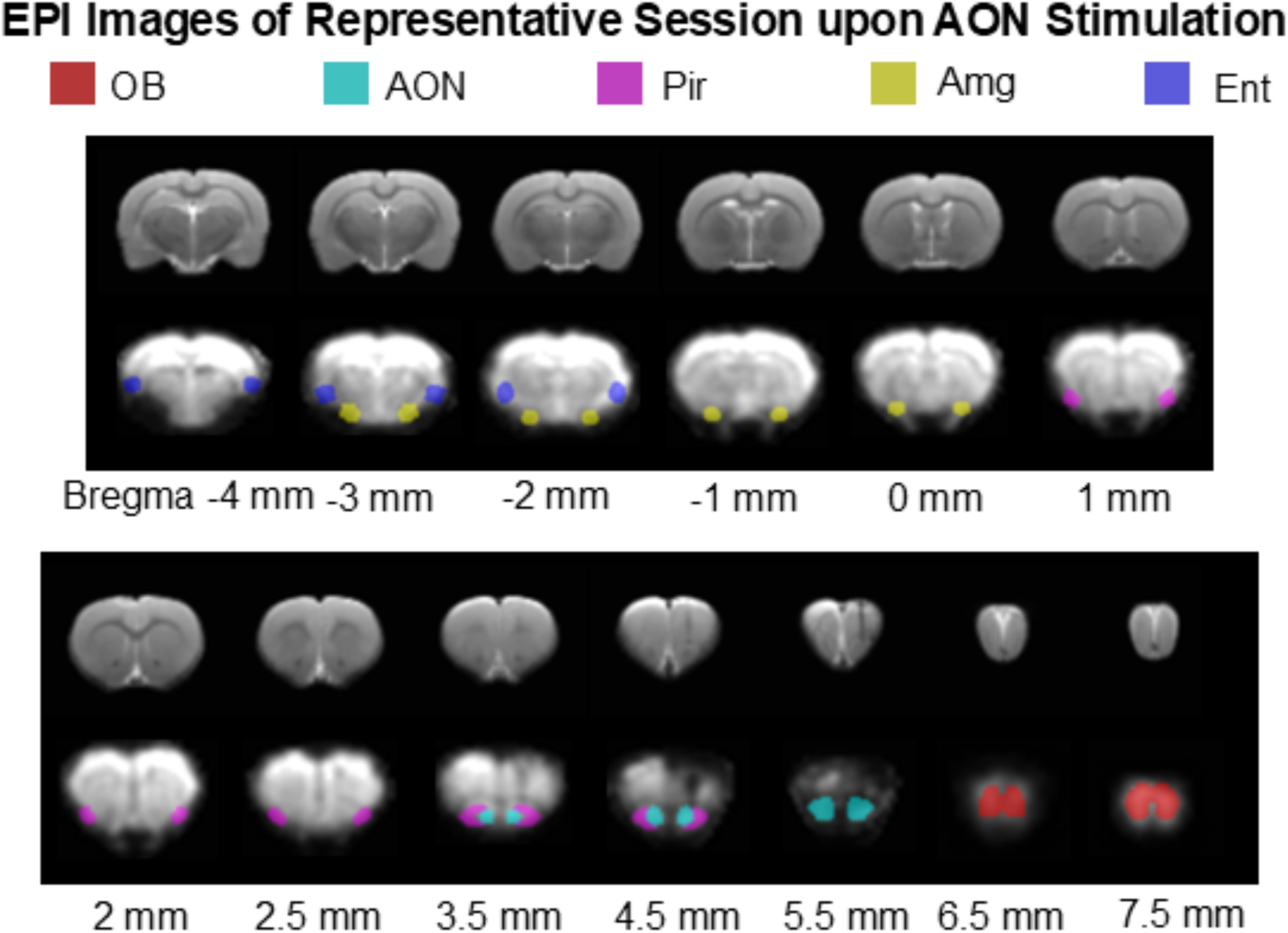
MRI images from a representative animal that underwent the AON optogenetic stimulation experiment. T2 images were displayed with their corresponding EPI images. ROIs including OB, AON, Pir, Amg and Ent are indicated with their respective colored overlays on the EPI images.

**Supplementary Figure 3.**
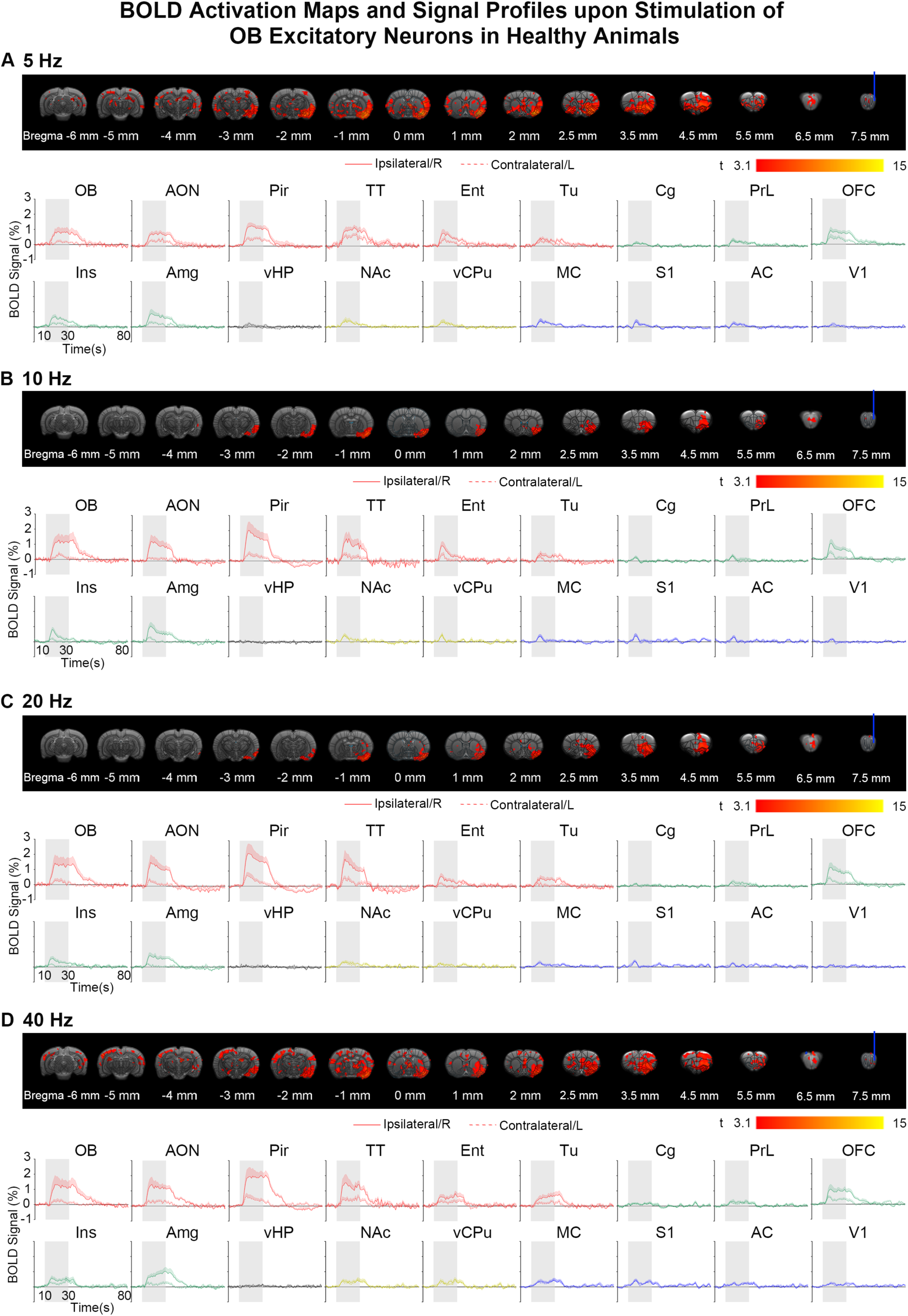
BOLD activation maps and BOLD signal profiles upon 5/10/20/40 Hz stimulation of OB excitatory neurons. **(A)** Bilateral activations in regions of the primary olfactory network and the corresponding BOLD signal profiles evoked by 5 Hz stimulation (n = 11; t > 3.1 corresponding to *P* < 0.001; error bars indicate ± SEM). **(B-D)** Localized activations in the ipsilateral hemisphere for regions in the primary olfactory network and the corresponding BOLD signal profiles evoked by 10/20/40 Hz stimulations (n = 11; t > 3.1 corresponding to *P* < 0.001; error bars indicate ± SEM). Note that the BOLD activation maps displayed in A-D were further corrected for multiple comparisons with threshold-free cluster enhancement with family-wise error rate (TFCE-FWE) at *P* < 0.05. *Abbreviations for the ROIs are as follows. Regions in the primary olfactory network: olfactory bulb (OB), anterior olfactory nucleus (AON), piriform cortex (Pir), tenia tecta (TT), entorhinal cortex (Ent), olfactory tubercle (Tu); limbic network: cingulate (Cg), orbitofrontal cortex (OFC), insular (Ins), amygdala (Amg); hippocampal network: ventral hippocampus (vHP); striatal network: nucleus accumbens (NAc), ventral caudate putamen (vCPu); sensorimotor network: motor (MC), somatosensory (S1), auditory (AC), visual (V1) cortex.*

**Supplementary Figure 4.**
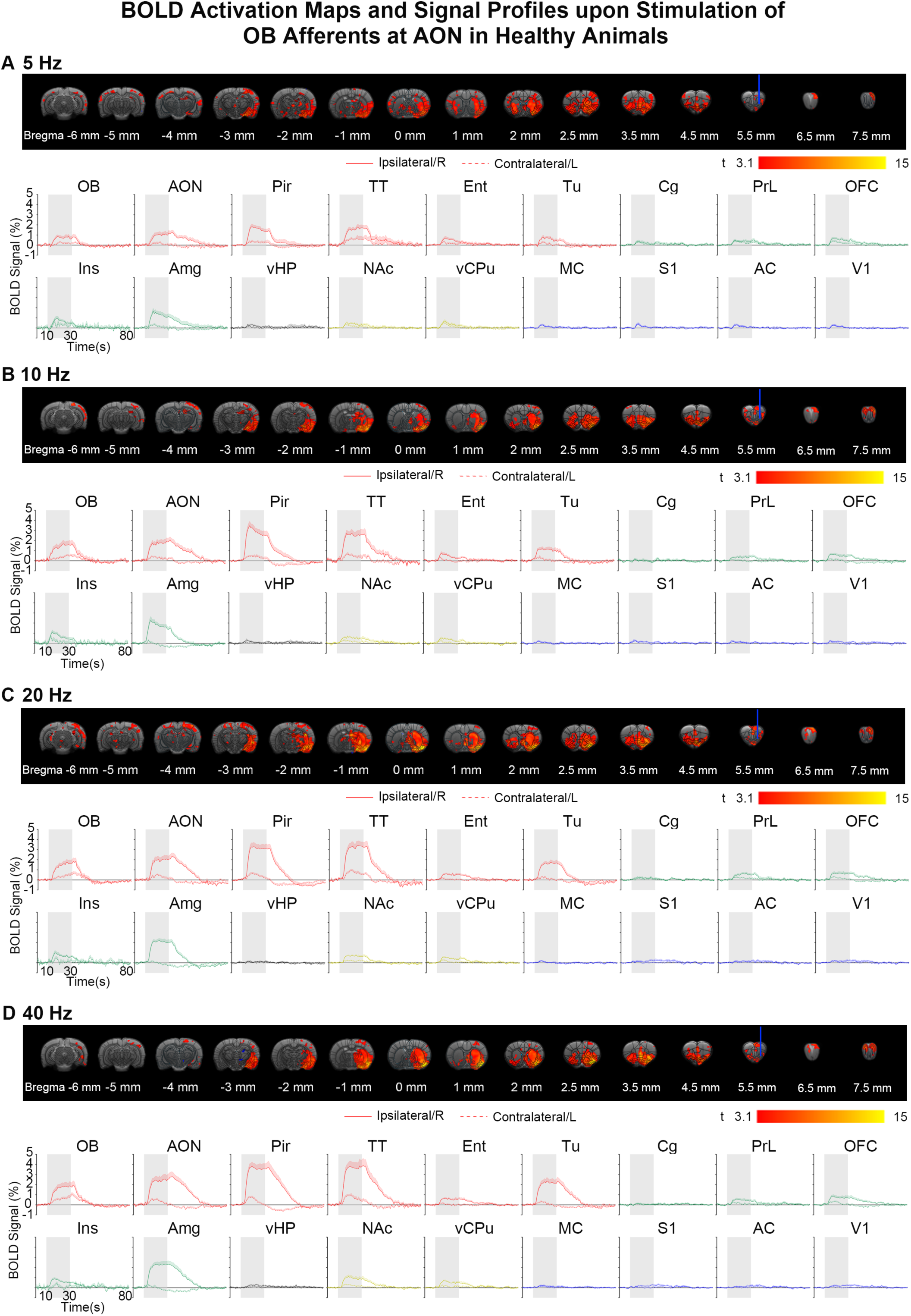
BOLD activation maps and BOLD signal profiles upon 5/10/20/40 Hz stimulation of OB afferents at AON. (**A-D**) Bilateral activations in regions of the primary olfactory, limbic and striatal network and corresponding BOLD signal profiles evoked by 5/10/20/40 Hz AON stimulations (n = 11; t > 3.1 corresponding to *P* < 0.001; error bars indicate ± SEM). Note that the BOLD activation maps displayed in A-D were further corrected for multiple comparisons with TFCE-FWE at *P* < 0.05.

**Supplementary Figure 5.**
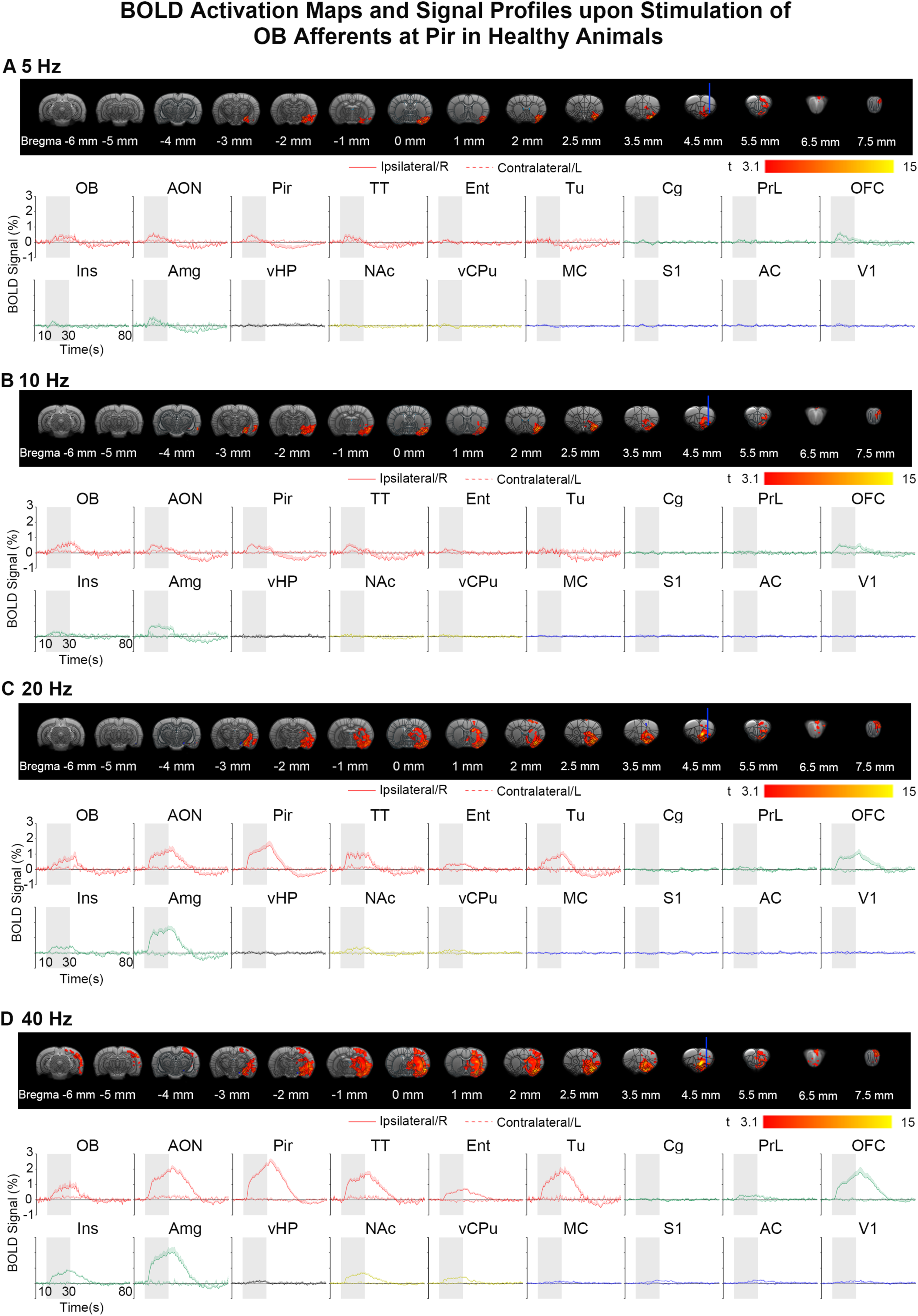
BOLD activation maps and BOLD signal profiles upon 5/10/20/40 Hz stimulation of OB afferents at Pir. (**A**-**D**) Localized BOLD activations in the ipsilateral hemisphere for regions of the primary olfactory network and the corresponding BOLD signal profiles evoked by 5, 10, 20, and 40 Hz Pir stimulations (n = 9; t > 3.1 corresponding to *P* < 0.001; error bars indicate ± SEM). Note that the BOLD activation maps displayed in A-D were further corrected for multiple comparisons with TFCE-FWE at *P* < 0.05.

**Supplementary Figure 6.**
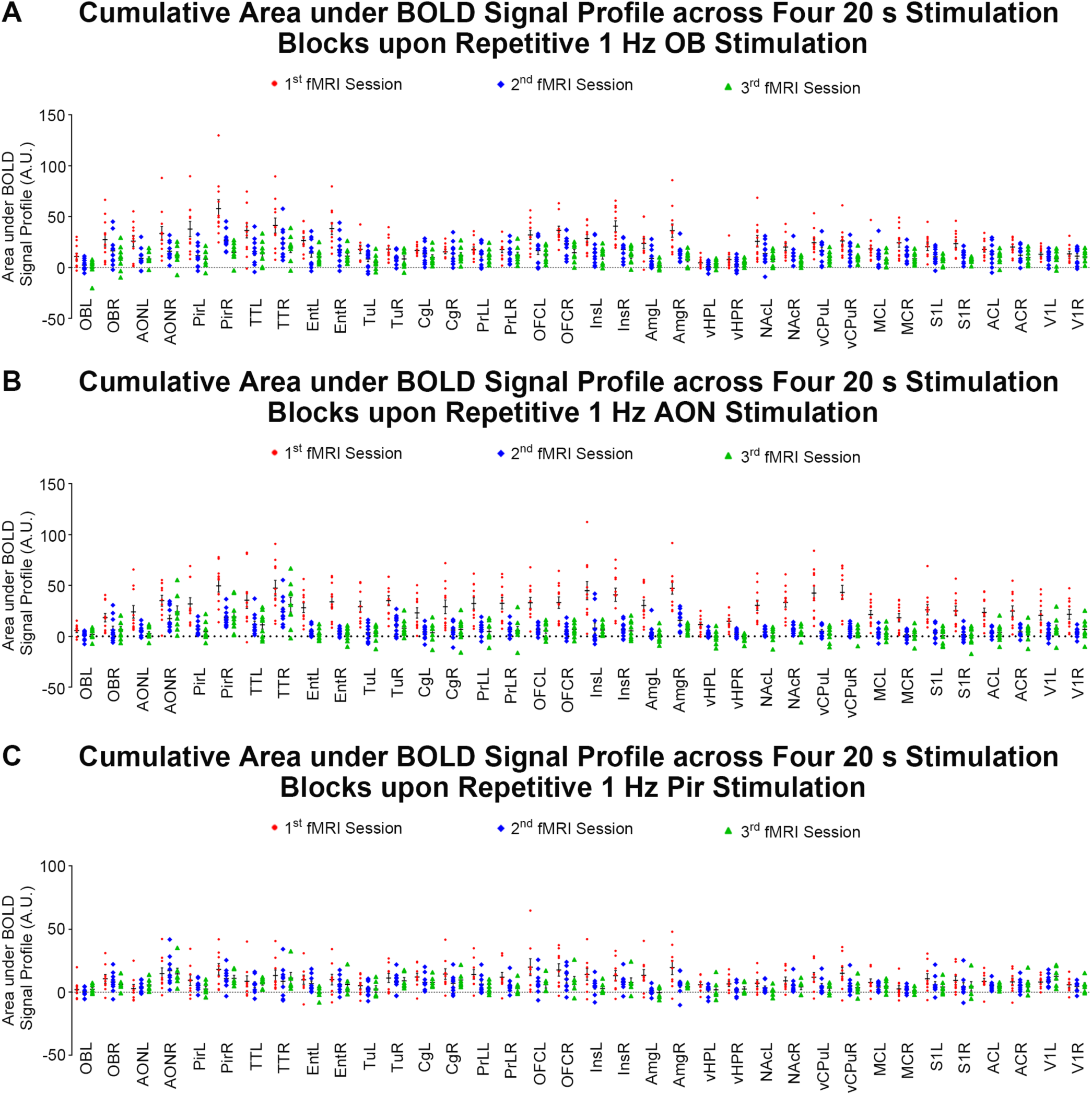
Repetitive optogenetic excitation of OB excitatory neurons and OB afferents at AON significantly decreased BOLD activations in olfactory networks. Summary of the cumulative area under BOLD signal profile over four 20 s stimulation blocks for each atlas defined ROIs in optogenetic fMRI experiments stimulating (**A**) OB excitatory neurons (n = 11), (**B**) OB afferents at AON (n = 11) and (**C**) OB afferents at Pir (n = 9) at 1 Hz. Note that all animal data points are shown. Red dots, blue rhombuses and green triangles represent the cumulative area under the BOLD signal profile computed for the 1^st^, 2^nd^ and 3^rd^ fMRI sessions, respectively. Error bars indicate ± SEM.

**Supplementary Figure 7.**
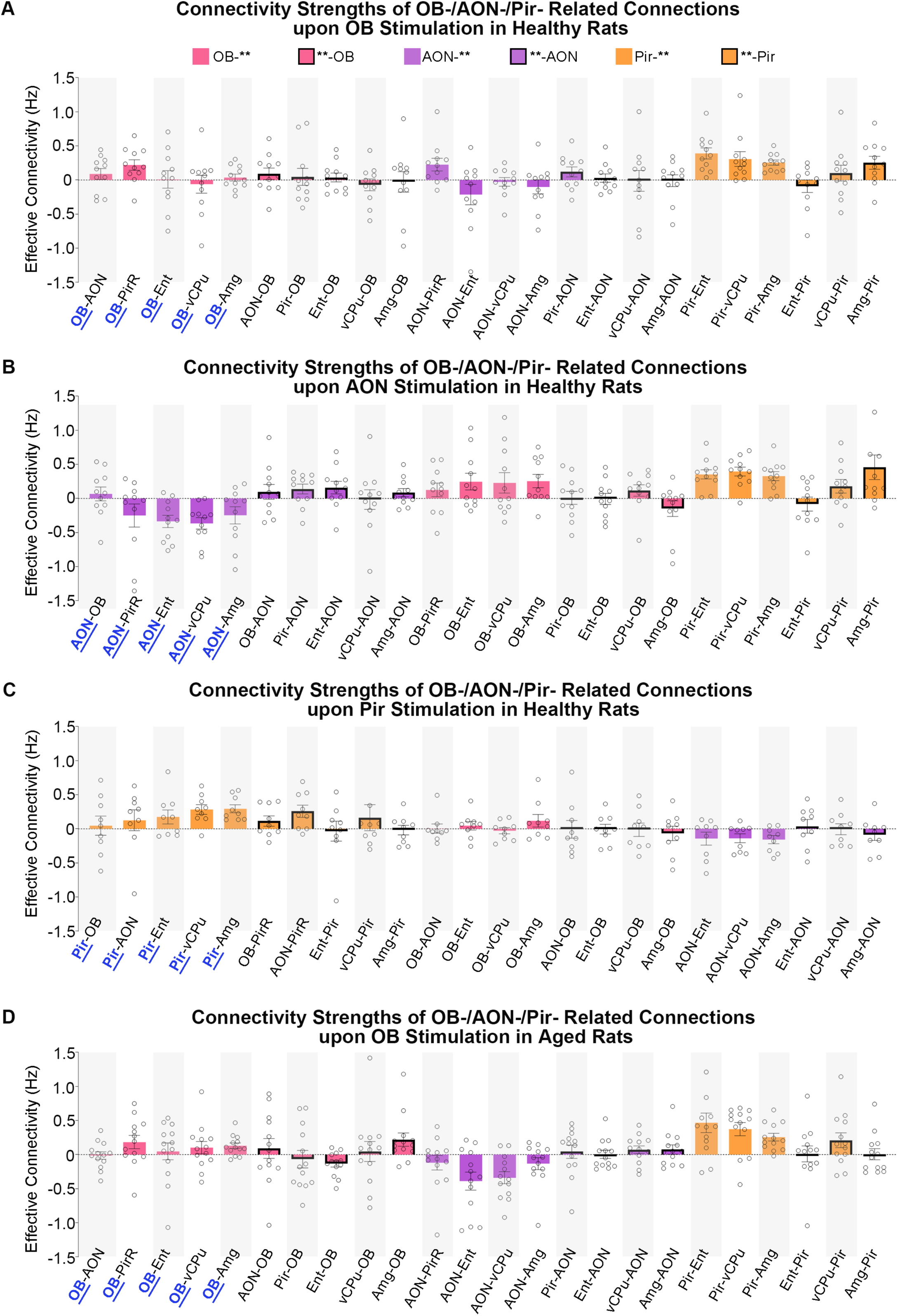
Summary of the effective connectivities in the optogenetic fMRI data that are modeled with dynamic causal modeling (DCM). Quantitative summary of the computed connectivity strength for each connection defined in the six-node olfactory network for DCM of 1 Hz optogenetic fMRI experiments stimulating (**A**) OB excitatory neurons (n = 11), (**B**) OB afferents at AON (n = 11), (**C**) OB afferents at Pir (n = 9), and (**D**) OB excitatory neurons in an aged rat model. The six-node olfactory network, which is described in Fig. 4A, encompasses regions from the primary olfactory (OB, AON, Pir and entorhinal cortex, Ent), striatal (ventral caudate putamen, vCPu) and limbic (amygdala, Amg) networks. Individual animal data points are shown and error bars indicate ± SEM.

**Supplementary Figure 8.**
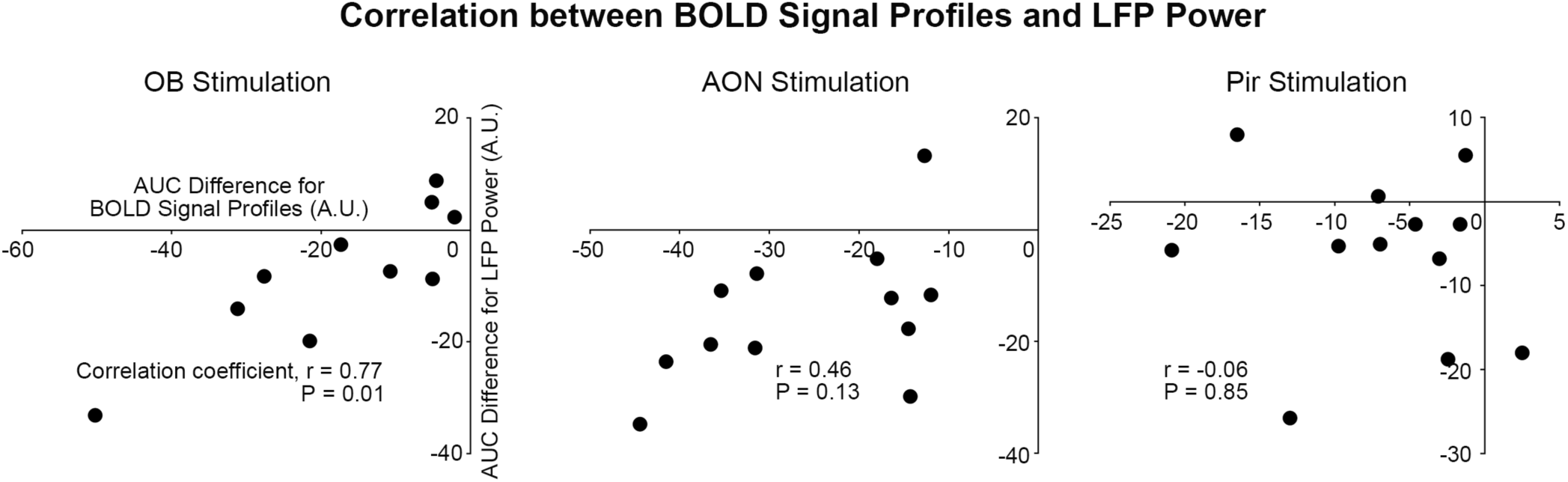
Correlation coefficient analysis between BOLD signal profiles and LFP power upon 1 Hz stimulation using Spearman nonparametric correlation. Relationship between the differences of LFP power and BOLD signal profiles showing adaptation were described by correlation coefficient, r and the corresponding P value. Differences in the AUC of BOLD signal profiles or LFP power were calculated between the 2^nd^ stimulation session vs. 1^st^ stimulation session. The r and P values for optogenetic stimulation of the three respective primary olfactory regions are OB: r = 0.77 (P = 0.01), AON: r = 0.46 (P = 0.13), and Pir: r = - 0.06 (P = 0.85).

**Supplementary Figure 9.**
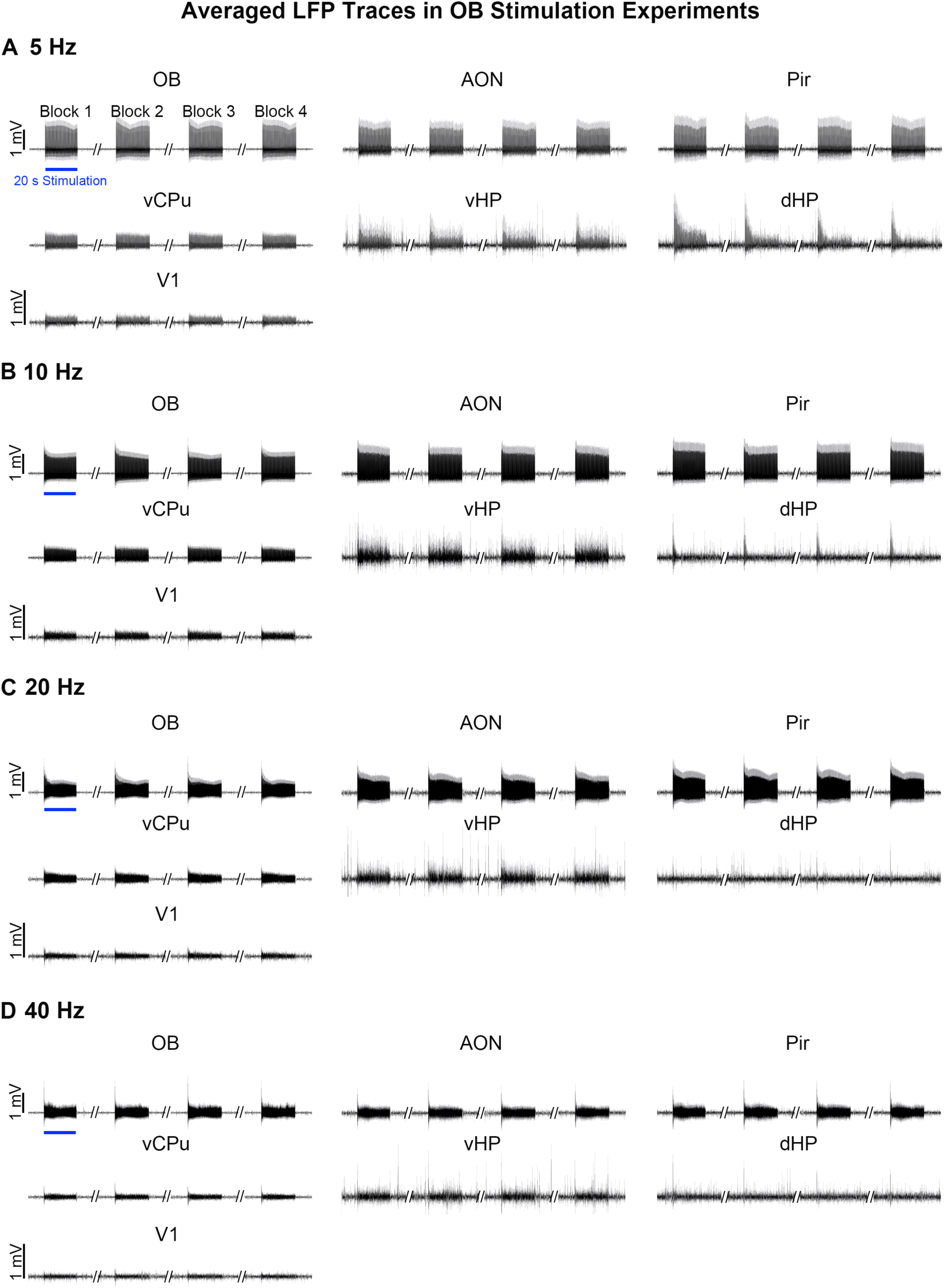
Local field potentials (LFPs) evoked by optogenetic stimulation of OB excitatory neurons at 5, 10, 20 and 40 Hz. (**A-D**) Averaged LFP traces of recordings at the ipsilateral OB (n = 7), AON (n = 7), Pir (n = 7), vCPu (n = 7), V1 (n = 7), vHP (n = 2) and dHP (n = 3) upon 5, 10, 20 and 40 Hz stimulations. Error bars indicate ± SEM.

**Supplementary Figure 10.**
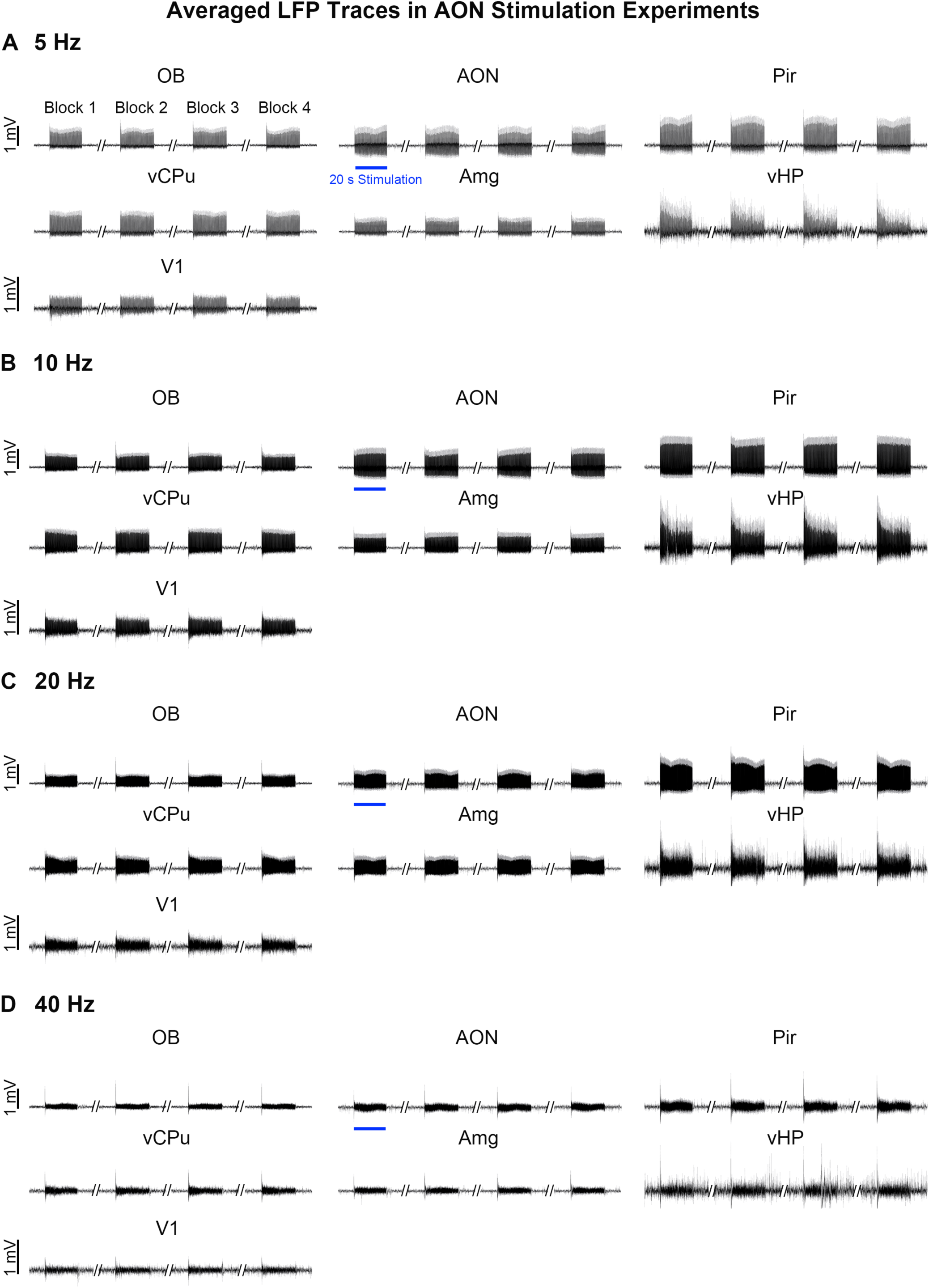
LFPs evoked by optogenetic stimulation of OB afferents at AON at 5, 10, 20 and 40 Hz. (**A-D**) Averaged LFP traces of recordings at the ipsilateral AON (n = 7), OB (n = 6), Pir (n = 5), vCPu (n = 3), V1 (n = 3), Amg (n = 4) and vHP (n = 5) upon 5, 10, 20 and 40 Hz stimulations. Error bars indicate ± SEM.

**Supplementary Figure 11.**
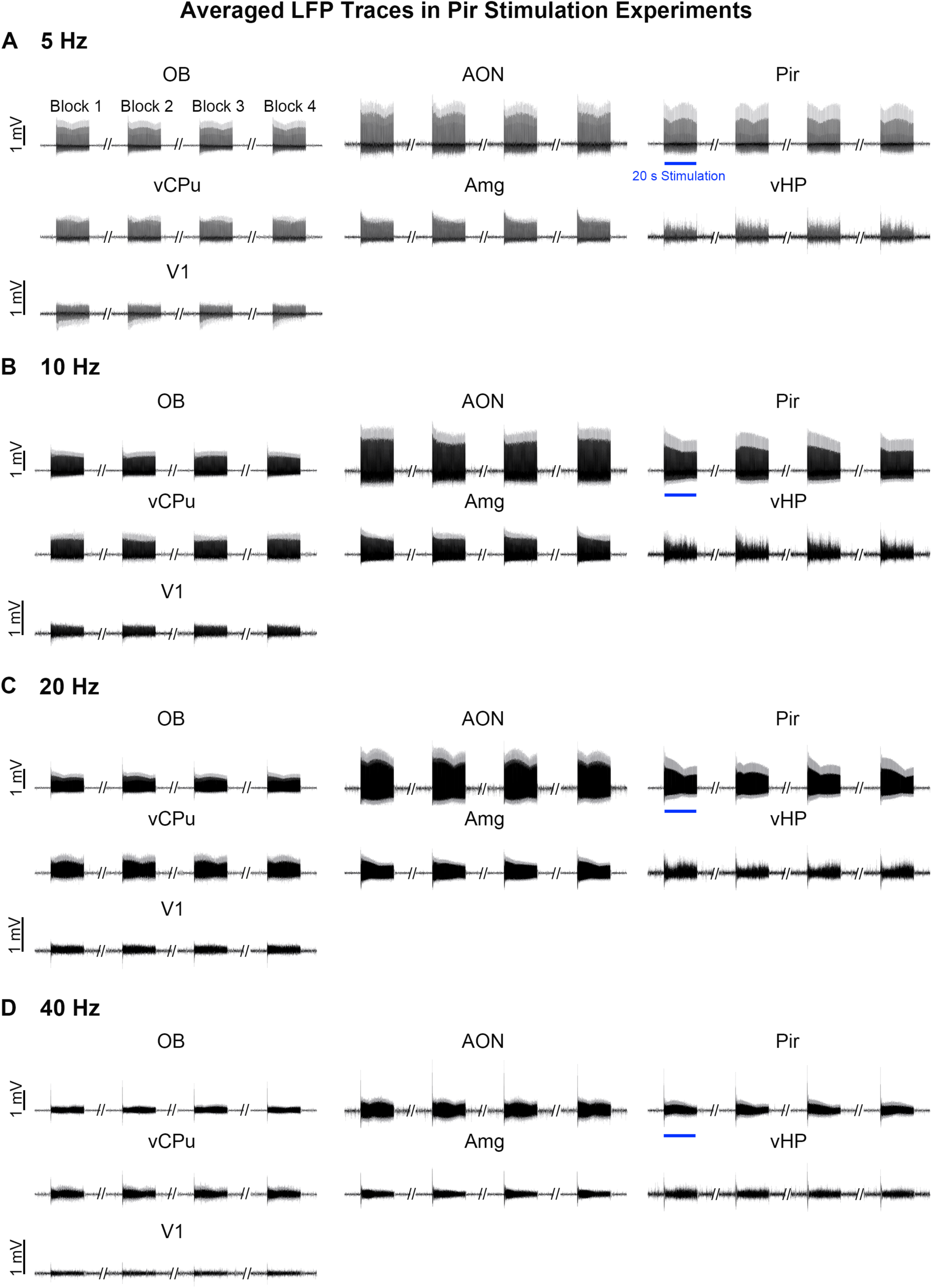
LFPs evoked by optogenetic stimulation of OB afferents at Pir at 5, 10, 20, and 40 Hz. (**A-D**) Averaged LFP traces of recordings at the ipsilateral Pir (n = 5), OB (n = 5), AON (n = 4), vCPu (n = 3), V1 (n = 3), Amg (n = 2) and vHP (n = 5) upon 5, 10, 20 and 40 Hz OB stimulations. Error bars indicate ± SEM.

**Supplementary Figure 12.**
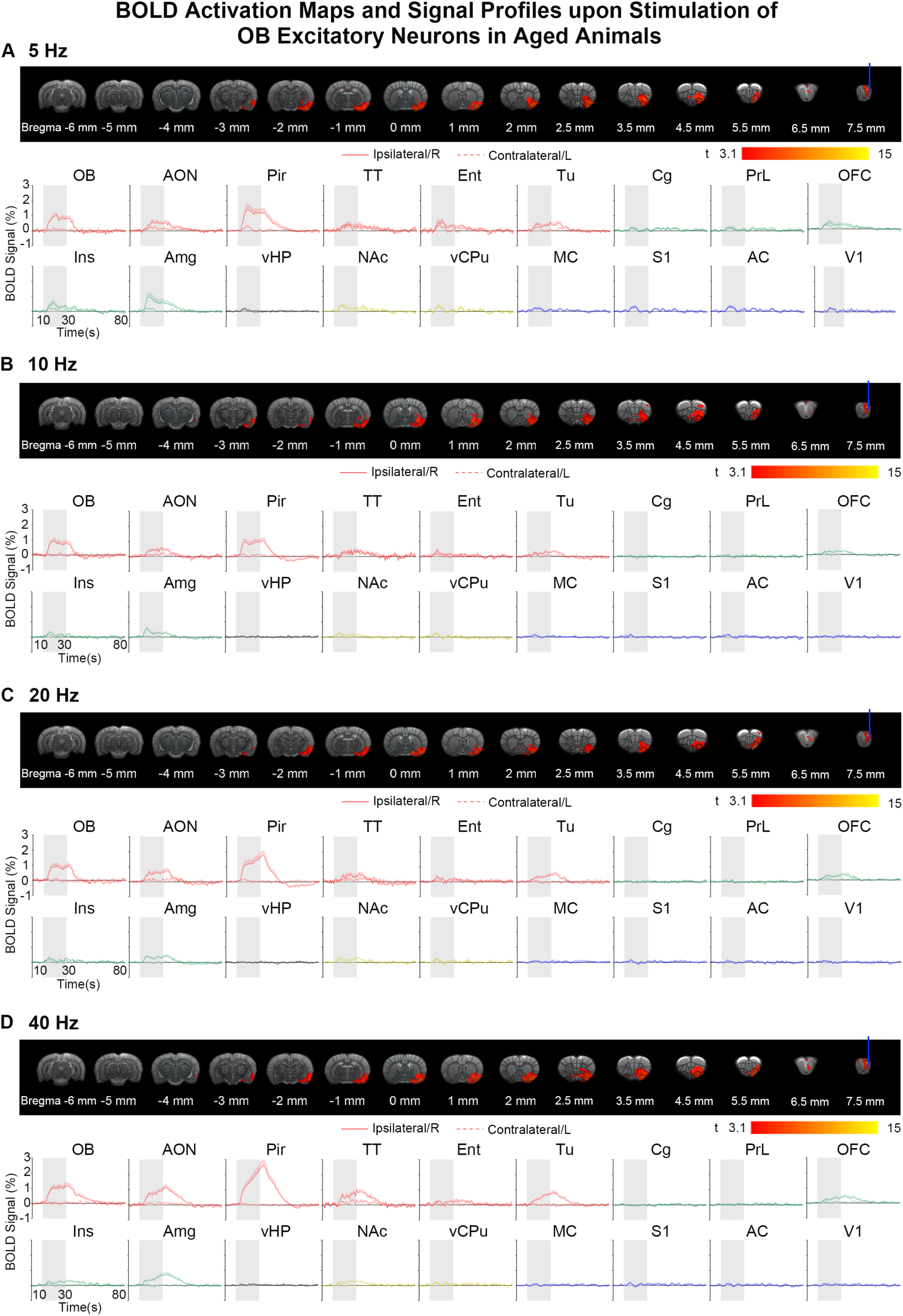
BOLD activation maps and BOLD signal profiles upon 5/10/20/40 Hz stimulation of OB excitatory neurons in an aged animal model. (**A-D**) Localized activations in the ipsilateral hemisphere for regions in the primary olfactory network and the corresponding BOLD signal profiles evoked by 5/10/20/40 Hz stimulations (n = 13; t > 3.1 corresponding to *P* < 0.001; error bars indicate ± SEM). Note that the BOLD activation maps displayed in A-D were further corrected for multiple comparisons with TFCE-FWE at *P* < 0.05.

